# “*Visualize, describe, compare*” – nanoinformatics approaches for material-omics

**DOI:** 10.1101/2025.03.27.645714

**Authors:** Cristina Izquierdo-Lozano, Marrit M.E. Tholen, Valentina Girola, Anna Świetlikowska, Maarten Merkx, Francesca Grisoni, Lorenzo Albertazzi

**Author notes:** Authors contributed equally (Cristina Izquierdo-Lozano, Marrit M.E. Tholen).

## Abstract

Bioinformatics and cheminformatics are established disciplines, but nanoinformatics, the development of computational tools for understanding and designing nanomaterials, is still in its infancy. In light of the new data-driven approaches for nanomaterials discovery, there is a growing need for in silico tools tailored to analyze nanomaterials datasets. This is particularly crucial for soft materials, where a crystalline structure cannot be obtained and therefore the characterization datasets are less structured, and there are no standard methods for data mining. Here we present a computational package capable of visualizing, describing, and comparing nanoparticle datasets obtained with super-resolution microscopy at the single-particle and single-molecule level. Our method allows us to: i) visualize multiparametric nanoparticle datasets to grasp material properties and heterogeneity; ii) have a quantitative evaluation of a material through a series of molecular descriptors, and iii) compare different materials quantitatively and globally, going beyond comparison of a single property.

We applied this method to a library of targeted nanoparticles revealing particle heterogeneity, similarities, and correlations between the synthesis and the physicochemical properties of the different nanomaterials. Finally, we show the potential of this approach to reveal batch-to-batch variations in time and between users hidden in standard analysis.

## 1 Introduction

In the current landscape of scientific research, where large dataset-based data-driven approaches are becoming powerful, informatics has become a cornerstone of advancements in various disciplines [1]. In biology, thanks to advances in analysis techniques such as -omics approaches [2], large structured datasets of the proteome and transcriptome [3] have been generated, and bioinformatics techniques have allowed the full potential of these datasets. This resulted in unique initiatives such as the Human Protein Atlas [4] and the Cancer Genome Atlas [5, 6] allowing researchers to access this data and use it to advance understanding of biological mechanisms and design novel therapies [7, 8]. Similarly, chemistry and drug discovery are moving to high-throughput and data-driven approaches based on chemoinformatics. [9, 10, 11]. Initiatives like the chEMBL or BindingDB databases [12, 13, 14, 15], are aiding drug discovery, as well as understanding structure-activity relations. [16].

Looking at the impact of cheminformatics and bioinformatics, it is logical that a similar approach would be extremely beneficial for nanotechnology and nano-materials. Indeed the field of nanoinformatics, despite being in its infancy, has the promise of transforming the way we discover, understand, and apply nanomaterials [17]. Nanoinformatics can provide a data-driven framework to tackle the complex materials design space’s navigation and implement the most advanced machine learning and artificial intelligence tools for materials design.

However, the fields of nanotechnology and materials pose unique challenges to informatics approaches that must be tackled before a widespread application of a data-driven approach. First, the material design space, that is, the sum of all possible structures available, is enormous compared to the already wide chemical space of small molecules due to the many combinations of components and conditions possible [18, 19, 20]. Compared to cells, where the same building blocks like nucleic acids and proteins are the basis for all systems, materials are much more varied in their structures. Therefore, any standardization effort necessary for data-driven approaches is significantly more challenging. Moreover, unlike small molecules, nanostructures are not perfectly defined entities [21]. This has two main implications: i) their description is significantly more complex as they cannot be expressed by a molecular structure or a set of simple molecular descriptors; ii) the structures are heterogeneous with high variations within the batch (polydispersity) and between batches. For these reasons, the first nanoinformatics approaches focused on crystalline inorganic materials that can be coded by their lattice structure obtained with crystallography [22, 23], whereas approaches for soft materials, the most widely used in nanomedicine, are completely lacking. Therefore, it is crucial to have informative standardized datasets for soft materials and new computational approaches to analyze them to enable nanoinformatics in the fields of nanobiomaterials and soft matter.

Here, we define a nanoinformatics framework to visualize, describe, and compare nanoparticles for biomedical applications tackling several of the nanoinformatics challenges for soft materials. We showcase its application on an imaging dataset of nanoparticles designed for drug delivery, obtaining unique information about nanoparticle dispersity, similarity, and reproducibility in an omics fashion.

These methods harvest data from the latest developments in multiparametric characterization and imaging. Because a chemical or crystalline structure is not accessible for soft materials, the most promising way to create informative datasets is to measure all relevant properties such as size, shape, composition, physical properties, and chemical functionalities with single-particle sensitivity. Although this was impossible up to now, the last decade’s innovations in imaging allow us to tackle this challenge. First, multiplexed and correlative imaging allows us to measure multiple features of materials using different fluorescent probes and/or different imaging modalities [24, 25, 26, 27, 28, 29, 30, 31, 32]. This, in combination with new developments of probes, each tailored to measure specific physicochemical properties of materials (e.g. polarity, viscosity, surface ligands), allowed us to build datasets of materials parameters as descriptors. Finally, new methods such as super-resolution imaging entered the materials characterization arena, providing nanometer resolution, single-molecule sensitivity, and compatibility with multiplexing approaches. Here we harvest on multiplexed super-resolution imaging to analyze a library of nanomaterials and obtain twenty-three material features at the single-particle and single-molecule level. This dataset is comparable with single-cell omics, due to the single-particle level at which the analysis was performed and the multiple parameters analyzed.

While these new techniques open the way to a data-driven understanding of materials and their design, the datasets obtained pose a great challenge to data visualization and analysis, and there is currently no tool to study the enormous amount of information obtained from these techniques. Typically, nanomaterials are analyzed one at a time for a specific property (e.g. size distribution [33, 34]), a frequency histogram of such property is plotted, and the polydispersity index (PDI) is calculated. When studying large datasets of tens of particles analyzed for many parameters, this approach does not make use of all the information contained in the data. Moving from a single-parameter analysis of properties and heterogeneity to a comprehensive multidimensional analysis will provide a holistic understanding of the nanostructures. This will also contribute to having potential relevance in material science, comparable to what omics techniques did in biology (material-omics). Note that established bioinformatics approaches cannot be applied directly due to the large differences in the nature of the data. Materials are extremely heterogeneous, often showing strong overlap between formulations, reproducibility between users is low, and time is often a problem.

Here, we provide a quantitative framework for data analysis and visualization tailored for nanomaterial development. We tackle this issue from three sides: i) we tailor visualization tools for multivariate single-particle datasets; ii) we define quantitative molecular descriptors for nanoparticles; and iii) we propose a framework for materials holistic comparison, beyond looking at a single property at a time.

This approach opens up new opportunities for material understanding, discovery, and rational design of new formulations and supports reproducibility and quality control in nanotechnology.

## 2 Results

### 2.1 Dataset

We showcase our method by applying it to an imaging dataset of nanoparticles designed for drug delivery. The dataset contains information on a nanoparticle library composed of antibody-targeted silica nanoparticle formulations, characterized with PAINT super-resolution microscopy similar to what was previously done by our group [29]. The enhanced resolution of this technique allowed us to visualize individual nanoparticles and to measure structural features such as the size of the nanoparticle and their aspect ratio. Moreover, the multicolor ability and specific labeling enable the characterization of multiple functionalities on the nanoparticle surface. In this case, nanoparticles are decorated with antibodies for cell targeting, a typical strategy for drug delivery, and the fragment antibody binding (Fab) domains and the fragment crystallizable (Fc) domains are specifically labeled and quantified. This specific antibody-nanoparticle combination was chosen as a model system for our methodology because of its easy manufacturing and accessibility and interest in the nanomedicine community.

The super-resolution analysis of the proposed dataset contains information on single particles and single molecules, resulting in a rich dataset with a structure similar to the bioinformatics data of single cells (Figure 2 a).

**Figure 1:**
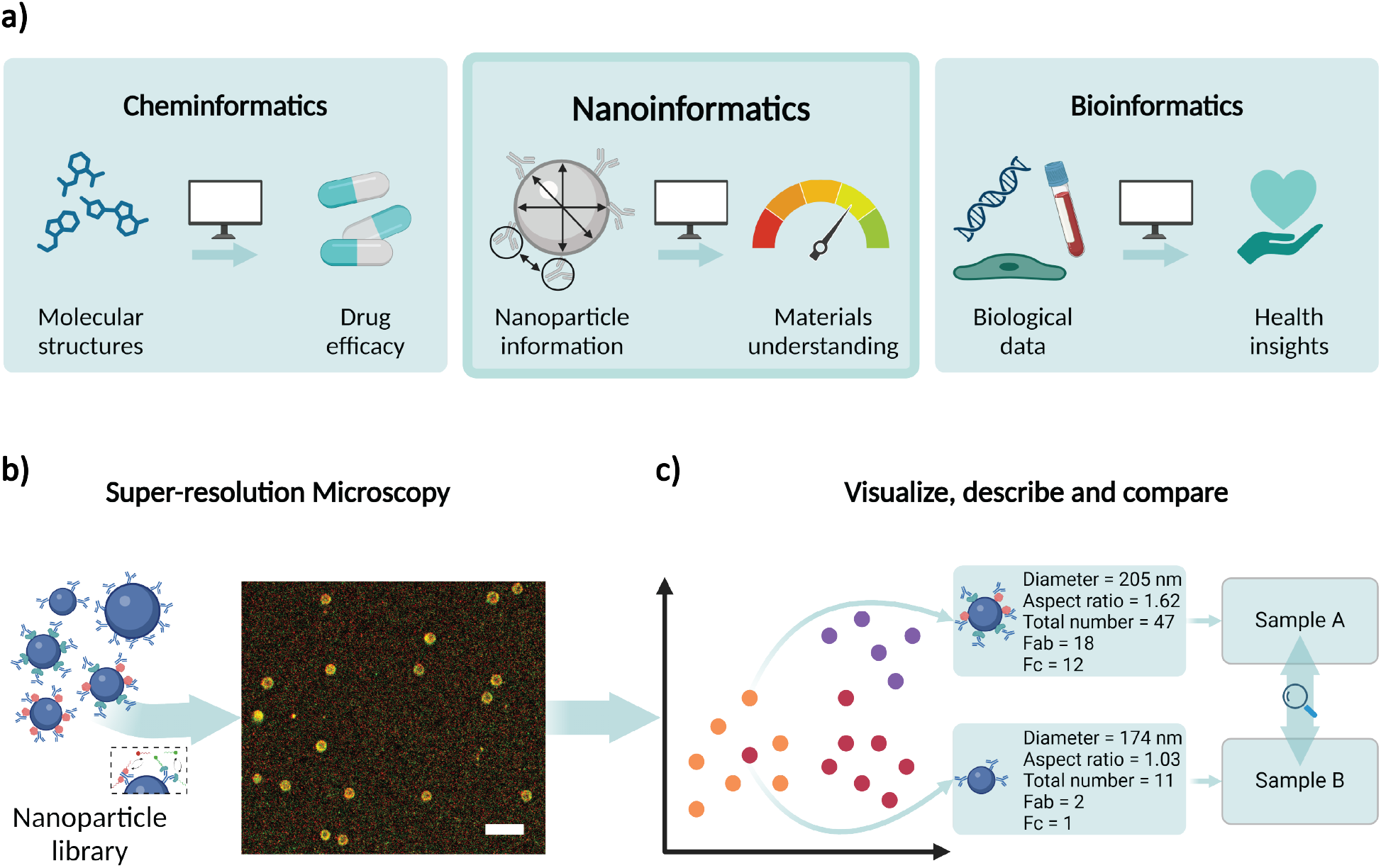
**a)** Main aspects and outputs of “Cheminformatics”, “Nanoinformatics” and “Bioinformatics”. **b-c)** Workflow to create and analyze our nanoparticle library. **b)** First, the different nanoparticle formulations are synthesized in the wet lab, after that they are labeled for imaging in super-resolution microscopy, and then the images are analyzed to extract the nanoparticle features, per single particle, **c)** to be finally used by machine learning methods to understand their relationship

**Figure 2:**
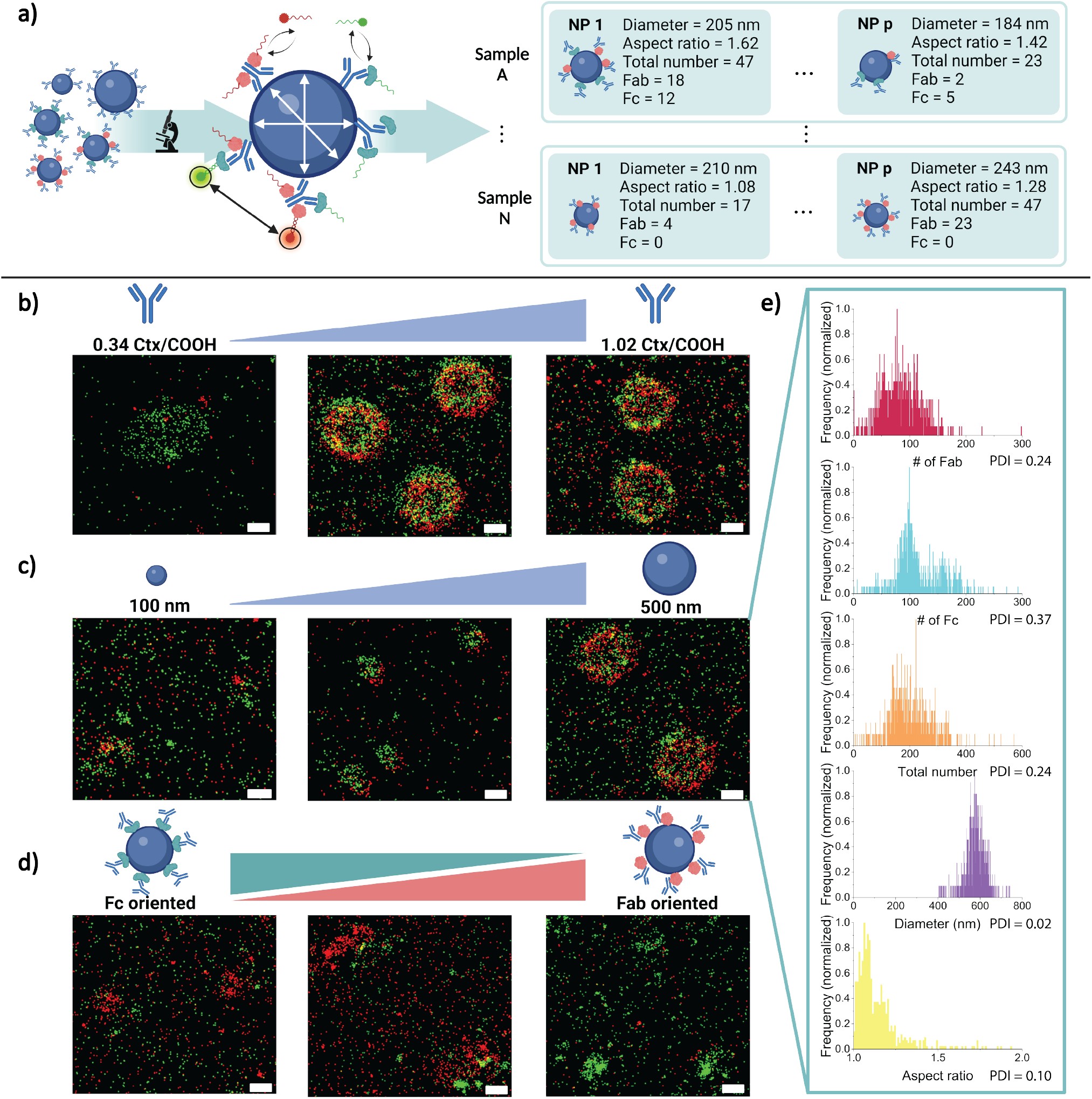
DNA-PAINT quantification of the number of exposed Fab and Fc domains on a nanoparticle library. Signal in red: Fc domains visualized by pG binding; signal in green: Fab domains visualized by pM binding. **a)** Workflow schematics. First, the library of nanoparticles is imaged using the DNA-PAINT method, then, the features of every single nanoparticle are extracted, thus building a dataset with 16 nanoparticle formulations, 20906 single-nanoparticle, and 23 features. **b-d)** Experimental variations to the different library formulations. **b)** nanoparticles were functionalized with a range of Cetuximab (Ctx) concentrations. Particles with a low amount of antibody per COOH group show a low number of localizations, while particles with a high number of Abs show a higher signal. **c)** nanoparticles of different diameters were functionalized with antibodies. **d)** nanoparticles were functionalized with a protein to obtain particles with a controlled antibody orientation. Different concentrations of the Fc-orienting protein pG (green) and Fab-orienting protein pM (red) were functionalized on the nanoparticle’s surface. This results in a higher signal of the other protein. **e)** Example of qPAINT results obtained with the *nanoFeatures* app

To show the potential of the analysis method, a nanoparticle library consisting of 16 particles was designed to vary systematically: i) size and morphology; ii) amount of antibody functionalities, and iii) conjugation strategy of the antibodies and therefore their orientation (Figure 2). The list of the diverse formulations can be found in the Supplementary Table S1.

Silica nanoparticles with different diameters ranging from 100 nm to 500 nm, encompassing various typical sizes used in nanomedicine, have been imaged and characterized (Figure 2c). The difference in sizes is visible in the images, which shows the ability of our microscope to resolve the nanoparticle structure. Silica nanoparticles with carboxylic acid (-COOH) functional groups were used to conjugate cetuximab antibodies (Ctx, a cancer therapeutic antibody targeting the EGFR biomarker) through EDC chemistry. To tune the valency of the nanoparticles, a key parameter for targeting, different ratios of Ctx/COOH were used: 0.34 Ctx/COOH (low), 0.68 Ctx/COOH (medium), or 1.02 Ctx/COOH (high) (Figure 2b). This variation in the amount of antibody can immediately be observed in the microscopy pictures, as the signal of both Fab (red) and Fc (green) domains increases with the amount of antibody added. The number of Fab-oriented and Fc-oriented antibodies are represented in our dataset as “target counts” for channels 1 (Fab) and 2 (Fc) and they are obtained from their signals by fitting a model on the bright and dark times of the nanoparticle, based on the qPAINT method [35, 36].

Finally, the library consisted of both oriented and randomly conjugated particles (Figure 2d). For oriented antibodies, amino-functionalized silica nanoparticles (NH2) were used, onto which proteins were immobilized for selective orientation (protein G and protein M). The antibody’s Fab domains and Fc domains were visualized using pM-DNA and pG-DNA respectively. These proteins are generic across species of antibodies but are specific to the domain. Due to the specificity of these proteins, the resulting nanoparticles preferentially expose one of the two domains of the antibody. In the microscopy images, it is shown that the Fc-oriented particles mainly show a signal in the red channel relating to the binding of protein G, while the Fab-oriented particles mainly show a signal in the green channel relating to the binding of protein M. To better represent the reality of nanoparticle formulations, the library was made by two researchers on multiple days, which adds to the heterogeneity of the formulations.

From these data, twenty-three features were extracted on a single-particle level using the *nanoFeatures* app [36]. This app utilizes a clustering algorithm (DBSCAN) for isolating the individual nanoparticles, after which the user can decide which features, e.g. number of localizations, diameter, and number of domains obtained via qPAINT, are reported. In Figure 2e, a selection of these reported features for a single sample is shown, showcasing the heterogeneous nature of the nanoparticle sample. The histograms in Figure 2e show the distribution of individual features such as size, aspect ratio, total number of Ab, and number of Fab / Fc domains. From this set of histograms, the complexity of the datasets is clear. All parameters are polydisperse, showing the presence of multiple populations in the same sample. While size and morphology are rather homogeneous, the amount of functionalities varies more wildly, resulting in a dataset where multiple heterogeneities of different levels overlap. Although plotting individual histograms provides useful information, this is far from perfect for understanding the data, and several questions remain unanswered. From this visualization and analysis, only the variation of single parameters is evaluated, but what is the global heterogeneity of the formulation? What is the correlation between features? Who is who on these histograms? - i.e. where a single nanoparticle lies in the different distributions? How to holistically compare two samples or the same sample replicated by two users?

To answer these questions, we provide a computational nanoinformatics framework for data visualization, description, and comparison in the following sections.

### 2.2 “Visualize”

The complexity of our multiparametric data presents a challenge in analysis and comprehension, as is the case in all high-throughput experimental methods, such as -omics approaches [37, 38] and necessitates the aid of visualization strategies that preserve the detail and information, but allow a user to understand it intuitively [39]. Even in the more established field of bioinformatics, there is an active discussion on the best way to visualize such complex datasets, such as single-cell gene sequencing, minimizing distortions [40, 41]. Ideally, the multidimensional data is represented in a 2D manner, in which the information about similarity is preserved, so two neighbors in the visualization also indicate relatedness [42]. In multivariate data analysis, many algorithms exist that allow us to visualize multi-variate data in lower dimensions. These would aid in understanding the heterogeneity and/or similarity, both within and between diverse nanoparticle formulations. As this is unexplored in the materials science field, in this work, we apply some of the most established techniques to the visualization of multivariate nanoparticle datasets, namely:

- *Principal Component Analysis* (PCA) [43, 44, 45]. Given a multivariate dataset, PCA identifies the main directions (principal components, or PCs) of data variation. Each PC is a new variable formed by a linear combination of the original variables. The PCs are as many as the original variables and are orthogonal (independent) from one another. The first PC captures the direction of greatest variance, the second PC captures the second most variance, and so on. By projecting the data onto these PCs, PCA reduces the dataset’s complexity while retaining the most significant patterns. Additionally, the linear coefficients (called loadings) of the original variables in each PC help interpret the visualization, providing insight into how the original variables contribute to the overall data structure.
- *t-Distributed Stochastic Neighbor Embedding* (t-SNE) [46]. t-SNE is a nonlinear method to visualize high-dimensional data. t-SNE aims to produce a low-dimensional (*e*.*g*., 2D or 3D) representation, where similar data points in the original data (‘neighbors’) are projected close together and dissimilar points are located farther apart. The number of neighbors used to preserve the local relationships in the data is controlled by the perplexity; the lower the perplexity, the more important the local similarities are compared to global similarities (and *vice versa*).
- *Uniform Manifold Approximation and Projection* (UMAP) [47] UMAP is another nonlinear method that, like t-SNE, helps reduce the dimensions of the data for easier visualization. Unlike t-SNE, UMAP attempts to preserve both local and global structures within the data, meaning that it captures not only close relationships between points but also the broader organization of the data. UMAP is often faster and more scalable than t-SNE and can better maintain the larger data structure when projected into lower dimensions.
- *Minimum Spanning Tree* (MST) [48, 49]. MST visualizes data in the form of a graph. MST shows the relationships (graph edges) between data points (graph nodes) to i) minimize the total distance between them, and ii) minimize the number of edges. MST reveals the underlying structure and ‘connectivity’ within the data, highlighting relationships within the data, *e*.*g*.., via hierarchical or branching data structures.

All methods come with respective advantages and disadvantages, and it is crucial to understand which is more suitable for nanomaterial characterization data. PCA is well known, fast, and easily explainable, thanks to the information on variance explained by each PC, and how the original variables contribute to the new visualization in the PC space, making it ideal for rapidly finding which structural or functional property of nanoparticles is more relevant in the dataset. Often, however, extensive multivariate data might contain complex non-linear information, which would not be captured by linear PCA (although non-linear variants exist [50]). In such scenarios, t-SNE and UMAP might be more suited for complex data, due to their ability to capture nonlinear information. This aspect comes at the expense of i) interpretability since the observed patterns cannot be mathematically traced back to the original variables, and ii) a slower and more complex optimization, which requires tuning several parameters (*e*.*g*., to control the effect of local vs. global information in the final visualization). Moreover, they might introduce distortions, *e*.*g*., due to the challenges in compressing high-dimensional data into lower dimensions and accurately capturing global and local information simultaneously. Finally, like UMAP and t-SNE, MST can capture non-linear information, but it does not need any optimization. However, computing the MST visualization can become computationally expensive.

In what follows, we compare each visualization technique in the selected case study (Figure 3), aiming to showcase how these visualizations can yield insights into complex patterns present in nanoparticle datasets, and ultimately to provide general indications for materialomics.

**Figure 3:**
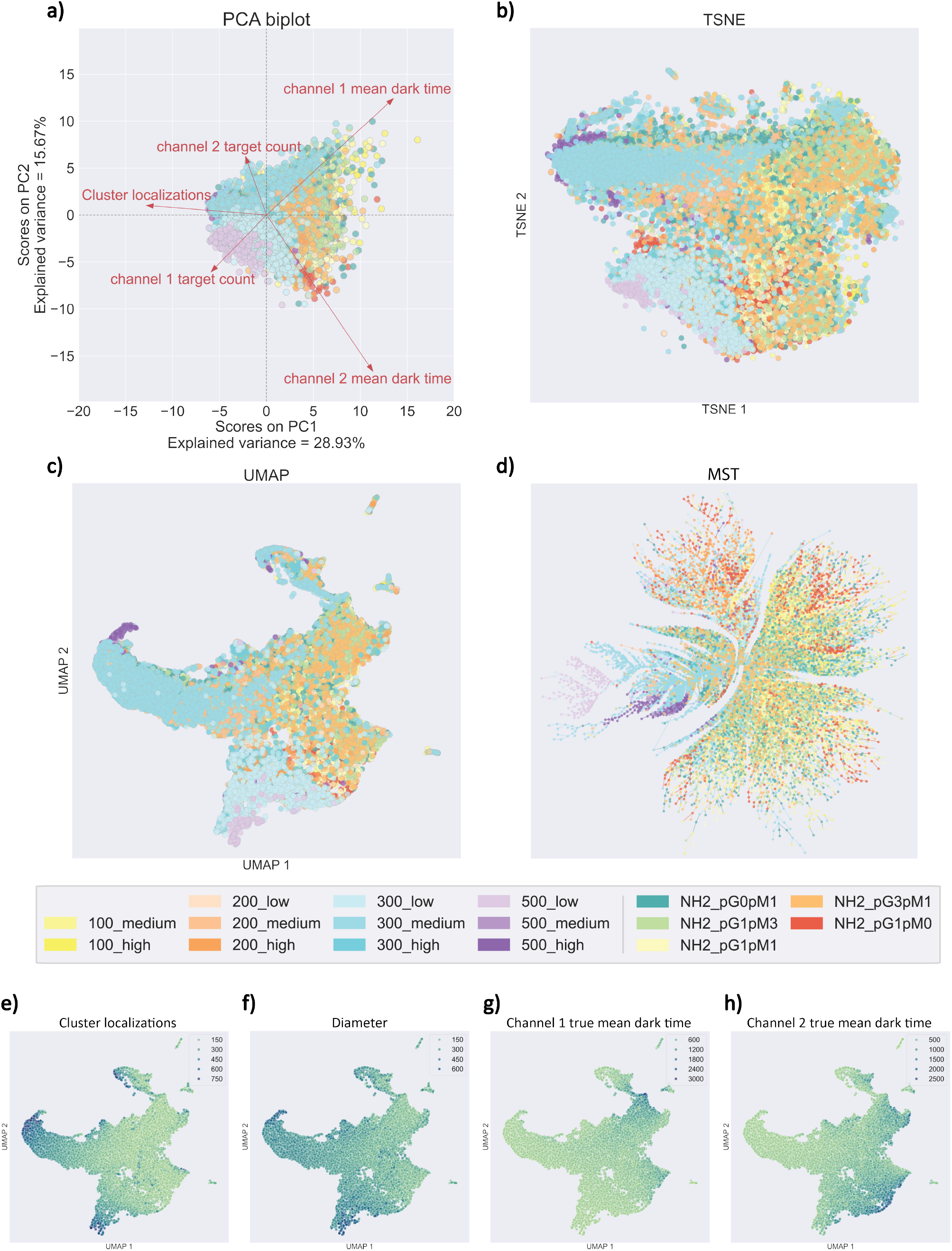
Visualization of Multiparametric Data, an overview. The 18 nanoparticles formulations, defined in Supplementary Table S1, are visualized in **a)** PCA, with the loadings overlayed on the samples (thus creating the ‘biplot’), **b)** t-SNE, UMAP, and **d)** MST. Finally, the UMAP plot was colored based on diverse selected features: **e)** Cluster localizations, **f)** Diameter, **g)** Channel 1 true mean dark time, which is the feature used to calculate the Fc count on the nanoparticle surfaces, and **h)** Channel 2 true mean dark time, which is the feature used to calculate the Fab count on the nanoparticle surfaces.

Visually inspecting all visualization outputs (Figure 3a-d) there are some common conclusions: i) the nanoparticle formulations tend to overlap and the samples do not appear as isolated clusters ii) nanoparticles within the same formulation tend to appear closer to each other (see Supplementary Figures S10-S18); iii) different samples seems to spread differently indicating different level of polydispersity. Overall, this indicates that each nanoparticle formulation has distinct features, making them occupy different portions of the feature space. However, because of the heterogeneity, they spread and partially overlap with each other, indicating that different formulations contain a fraction of particles of the same size and functionality.

When it comes to PCA, the first two principal components explain approximately 45% of the variance (Figure 3a). This variance is considered in general low and suggests that a remarkable portion of the data variability remains unexplained by the first two PCs and that additional components may be needed to capture a more comprehensive understanding of the dataset structure. For instance, four principal components would be needed to explain almost 70% of the variance (Supplementary Figure S9). Despite the low variance explained, the obtained PCA can be interpreted by looking at the variable coefficients (Figure 3a). When looking at the variability of the five most important variables (channel 1 mean dark time, channel 2 mean dark time, total cluster localizations, channel 1 target count, and channel 2 target count), we can see that the features relating to antibody orientation (mean dark time and target count per channel) differentiate the nanoparticle samples mostly along the second PC (y-axis, PC2), and features relating to the nanoparticle surface functionalization and size (cluster localizations) separate the nanoparticle samples in the first PC (x-axis, PC1). Thus, the features most influencing PCA variance are mainly tied to antibody functionalization and size, as diameter and antibody concentration are correlated and partially inferred from cluster localizations. This is expected since these parameters were chosen as the added variability in our library.

In the t-SNE and UMAP visualizations (Figure 3b and c), nanoparticles tend to group based on their formulation better than what was previously visible via a PCA (Supplementary Figures S12, S13, S14, and S15). Also, in this case, different formulations partially overlap, although to a lesser extent. This suggests that t-SNE and UMAP emphasize within-formulation similarity in addition to between-formulation similarity. This is most likely due to their ‘neighborhood’-based nature. Compared to t-SNE (Figures 3b), UMAP shows a better separation (Figure 3c) of different nanoparticle formulations, showing its ability to also capture global information in addition to local information.

Moreover, to improve UMAP explainability, we have colored the UMAP embeddings based on multiple features of interest (Figures 3e-h). In these plots, we can observe the gradient created by the values of these selected features and how they group the data into different areas of the plot. Therefore, we can relate the separation of the different nanoparticle formulations in Figure 3c to the specific feature or feature combination that caused it. Similarly to PCA, in these examples we see how the data is separated along the x-axis based on the nanoparticle’s antibody functionalization and size. Finally, in the MST visualization (Figure 3d) we can observe how nanoparticles of the same formulations are the leaves linked to the same common branch, which, at the same time, share the root with experimentally similar formulations. For example, the 500-nm formulations with three different levels of Ab functionalization are in the same root, but on different leaves. Moreover, we can observe that the formulations corresponding to the experiments with a controlled orientation of the antibodies (NH2 *) are found overlapping in the same branches as the 200 nm formulations, which is expected since the NH2 formulations are also formulated with 200 nm diameter nanoparticles, thus indicating that the diameter feature plays an important role in the cluster separation. This is potentially interesting as it would show the hierarchy and similarity between materials. For example, sets of particles with a common “ancestor” material. However, it is relevant to mention that a context is needed to avoid introducing potential biases and artifacts. [51]

This example shows how, in general, all visualization tools might lead to similar conclusions about general and broad patterns within the data. At the same time, each visualization approach has distinct advantages and disadvantages and might highlight different aspects of the underlying complexity. Overall, when explainability is the goal, PCA should be the go-to choice (given that sufficient variability is explained by the considered PCs). When visualization of nanoparticle (dis)similarities is needed, then UMAP is recommended, thanks to its ability to capture nonlinear information, its good trade-off between local and global information, its scalability, and its reduced computational requirements. Finally, in certain scenarios, MST could aid in highlighting hierarchical relationships, but it might come at the expense of computation time when large datasets are to be visualized.

### 2.3 “Describe”

Given the complexity of multivariate nanoparticle data, a proper quantitative description becomes key for characterizing their properties. To date, nanoparticles are often described in a univariate manner, *i*.*e*., by observing one variable at a time. For instance, the well-established polydispersity index (PDI, Eq. 2), is routinely used to quantify the size distribution of nanoparticles, focusing on a single parameter such as particle diameter. In this work, we extend ‘univariate’ measures of dispersion to multivariate nanoparticle data, hence extending their applicability to capture several aspects of nanoparticle datasets simultaneously. This global metric allows for the creation of an ‘identity card’ (ID card), where nanoparticles are characterized by a comprehensive set of parameters (Figure 4) for follow-up analysis.

**Figure 4:**
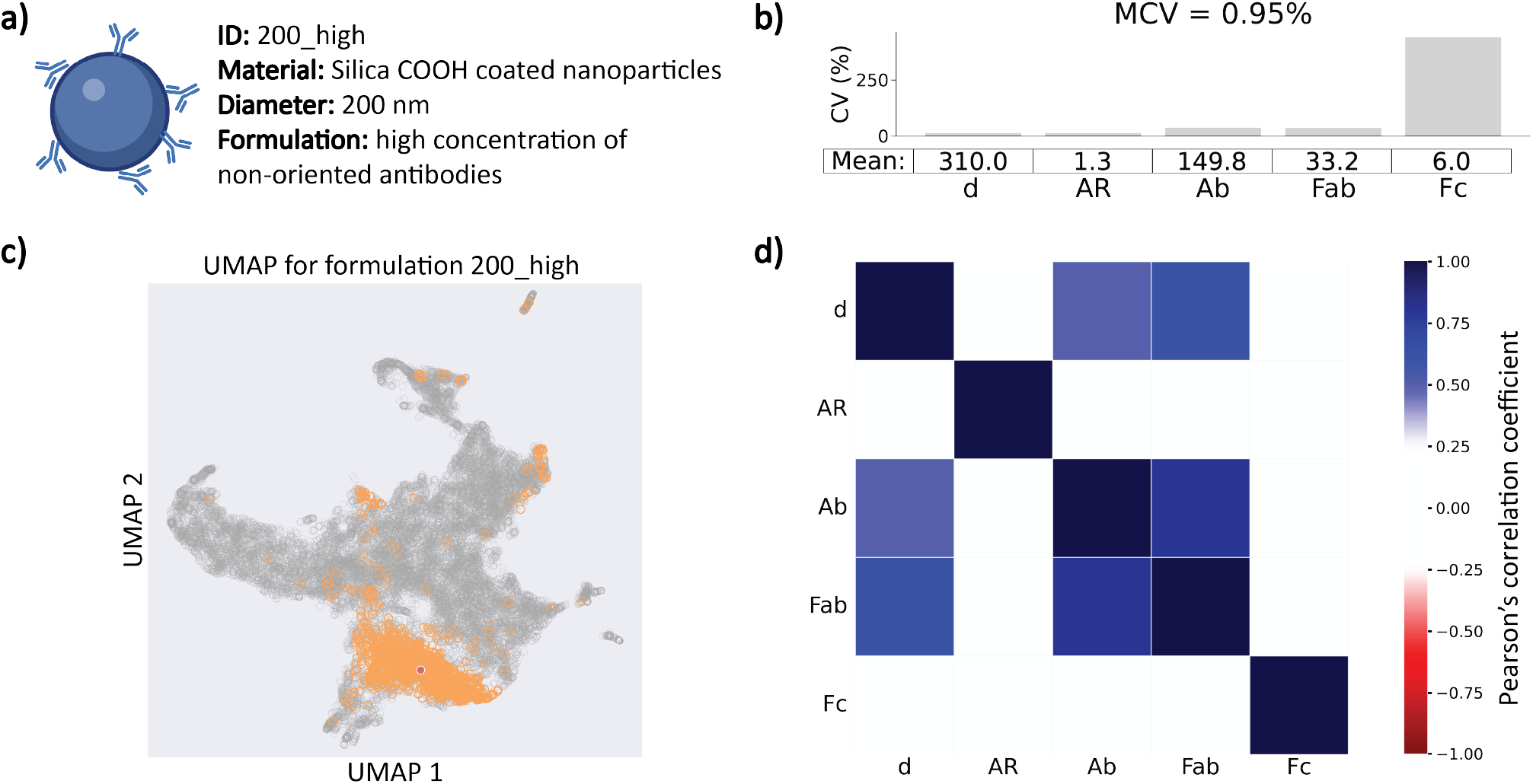
Example of a nanoparticle formulation ID card. In this case, we selected the 200 nm with high antibody concentration formulation and chose the ‘most-used’ nanoparticle features to show in the figures: Diameter (d), Aspect Ratio (AR), Cluster localizations, or antibody concentration, (Ab), channel 1 target count (Fab), and channel 2 target count (Fc). However, the calculations were done, per sample, using the complete dataset. **A)** Visual and text definition of the nanoparticle formulation. **B)** Individual CVs and mean values for each of the selected features, and MCV calculated using the multivariate space of the nanoparticle formulation. **C)** UMAP visualization highlighting the specific sample for this case study. The red dot indicates the center of mass of the nanoparticle formulation. **D)** Pearson’s correlation matrix, where blue indicates a direct correlation and red inverse correlation.

In this work, we collect several measures of ‘dispersion’ and the relationship of and between the features:

- *Feature correlation* (*r*), which captures the inter-dependency between different nanoparticle features, *e*.*g*., diameter, aspect ratio, etc. In this work, we calculate the pairwise Pearson’s correlation coefficient (Eq. 1), which ranges from −1 (perfect inverse correlation) to +1 (perfect correlation).
- *Coefficient of Variation* (CV). Given a feature of interest, the coefficient of variation measures the dispersion of its raw values around the average value (Eq. 3). This metric can be considered as an extension of the PDI to virtually any nanoparticle feature. CV ranges from 0 to infinity (the larger, the greater the variability of a feature).
- *Multivariate Coefficient of Variation* (MCV). The Multivariate Coefficient of Variation (MCV) [52] is an extension of CV to the case where multiple features must be considered simultaneously. The MCV ranges from 0 to infinite; the higher the MCV, the more diverse the features. This can be interpreted as a global PDI taking into account all nanoparticle features.

To showcase how these metrics can be used to describe the data, we have selected one formulation (nanoparticles with 200 nm diameter and high antibody concentration). The correlation matrix indicates how the features relate to each other. For example, in the selected case study (Figure 4d) the diameter and the total number of localizations are highly correlated (*r >* 0.50). This is a consequence of the fact that the larger a nanoparticle is, the higher the number of antibodies it hosts on its surface.

The CV captures the relative variability of the feature’s values compared to the average – the higher the CV, the more heterogeneous the values. In the selected example (Figure 4b), the diameter and aspect ratio are almost constant, as visible by their low CV (*CV* = 14.1 and *CV* = 14.3, respectively). On the contrary, the number of Fc-oriented antibodies has a higher relative variability of values compared to the average, as indicated by a higher CV value (*CV* = 441.1), suggesting that most of the heterogeneity of this nanoparticle formulation comes from Fc domain exposure. We observed as a general trend that surface ligands are more polydisperse in comparison to size and aspect ratio, indicating that synthetic procedures are more precise concerning structural features rather than functional ones.

Finally, the MCV can help understand the ‘global’ heterogeneity of the formulations, by capturing the variability in the multivariate space (*i*.*e*., by considering all features simultaneously, rather than just one or pairs at a time). The MCV has the advantage of being independent of the scale (*i*.*e*., measuring units) of the variables to be analyzed since it uses the covariance matrix to adjust for differences in feature variances (Eq. 4). To determine if the MCV is ‘high’ or ‘low’, it must be evaluated in context, as it depends on the range of the values given. In our nanoparticle library, the maximum is 57.74% and the mean is 10.80%. Therefore, in this case, the MCV is low (*MCV* = 0.95%), which denotes low dispersity within this nanoparticle formulation. This aligns with the low CV values seen before (Figure 4b) and the closeness of the samples from the same nanoparticle formulation in the UMAP space (Figure 4c).

These metrics and the visualization (Figure 4c) can be combined to form a ‘materials ID card’. Given a complex nanoparticle dataset, every formulation can be characterized by (a) uni- and multivariate metrics (*r*, CV and MCV), to gain insights into its dispersion, heterogeneity, and relationship between features, and (b) visualization (*e*.*g*. PCA and UMAP) as an indication of its positioning in the experimental space. The visualization plots can be further supported by the MCV value since they capture some overlapping information. For instance, the more spread a formulation is in the experimental space, the higher the MCV value. This can be observed by comparing the visualization plots, from any of the dimensionality reduction algorithms (Supplementary Figures S10-S15, to their corresponding plot containing their metrics and correlation matrix, for each nanoparticle formulation (Supplementary Figures S5 and S4).

We believe that this set of metrics achieves a good trade-off between information content and interpretability and might be suited to standardize the description of complex nanoparticle data across scientists and laboratories.

### 2.4 “Compare”

Finally, comparing two formulations can provide insight into their (dis)similarities, with various goals, *e*.*g*., deciding which formulation would be best for any specific application. However, to date, the comparison of diverse formulations is done based on one variable at a time (*e*.*g*., via histograms or boxplots).

Here, we introduce the Silhouette score (Eq. 5) as a way to compare nanoparticle formulations. Originally introduced to quantify the effectiveness of clustering [53], we applied it for the first time to quantify the (multivariate) overlap between two nanoparticle formulations. Given two formulations to be compared, a silhouette score of 1 indicates that the nanoparticles are highly similar within their formulation and distinct from the other formulation. A score of 0 suggests that the nanoparticles are at the boundary between the formulation groups, with overlapping formulations. A score of −1 implies that the nanoparticles fit better with a different group, indicating poor differentiation between formulations.

As a proof-of-concept, we showcase how the Silhouette scores – alongside the visualization and description approaches previously introduced – can be used to compare different formulations and analyze the same formulation prepared by different users or on different days.

First, we compare two pairs of formulations: (a) two oriented-antibody formulations (100% Fc-oriented antibodies, NH2 pG0pM1, and 100% Fab-oriented antibodies, NH2 pG1pM0) (Figure 5a-d), and (b) two non-oriented antibody formulations (100 medium and 300 medium) (Figure 5e-h).

**Figure 5:**
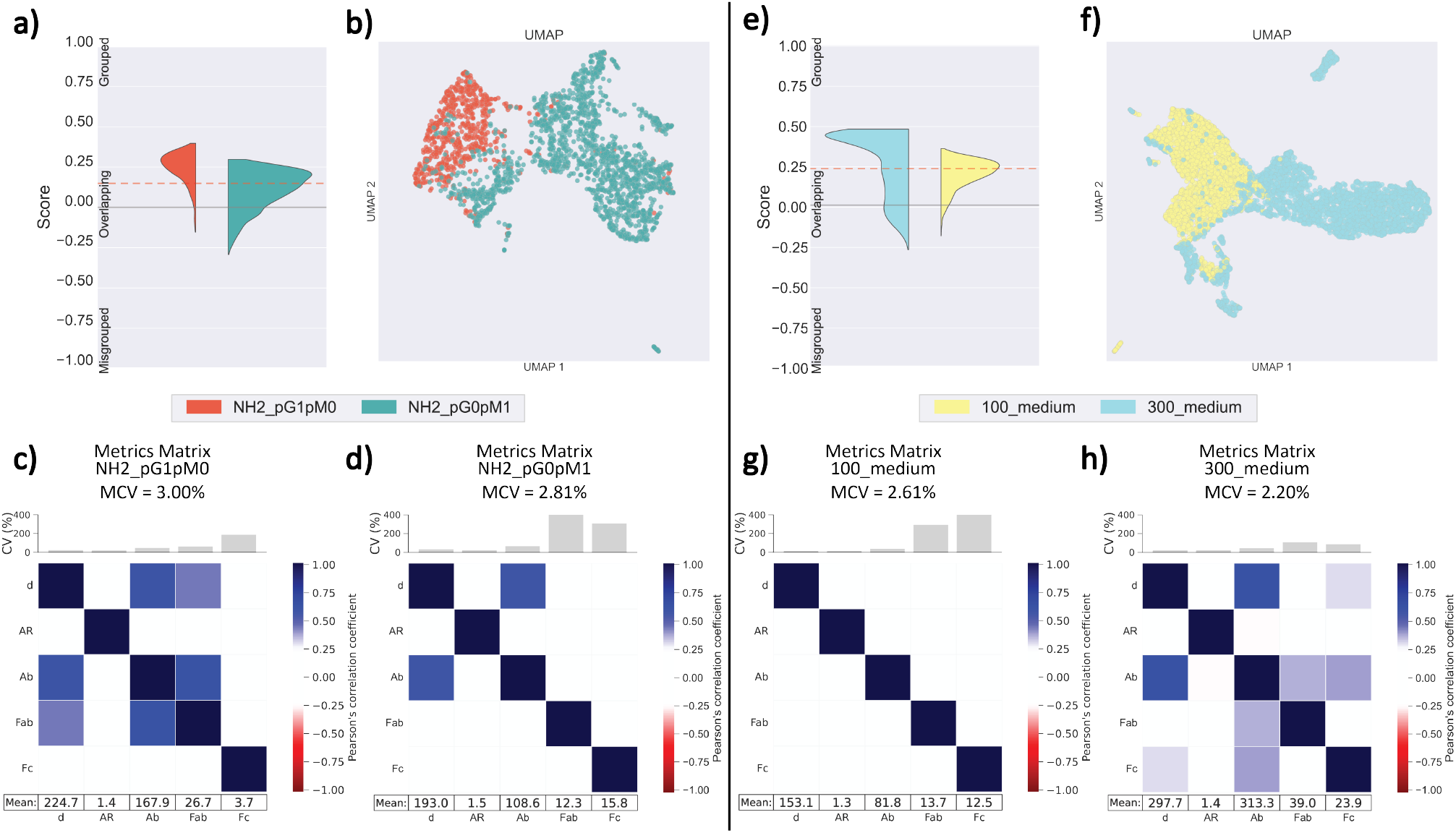
Pairwise comparison of two pairs of nanoparticle formulations. **a-d)** Oriented antibody formulations NH2 pG1pM0 and NH2 pG0pM1. **a)** Silhouette scores for the chosen pairs. The grey line marks the 0 value, and the red dotted line marks the average silhouette value between both formulations. **b)** UMAP plot recalculated for the chosen pair. **c-d)** Metrics matrices for the pair NH2 pG1pM0, showing MCV, CVs, and mean values for each selected feature. **e-h)** Non-oriented antibody formulations 100 medium and 300 medium.

The separation between the two pairs of formulations (as visible in the UMAP plots, Figure 5b and 5f) can be now quantified numerically by using the Sil-houette scores (Figures 5a and 5e). The samples from Fab-oriented antibodies formulation generally position closer to each other in the UMAP than samples from Fc-oriented antibodies formulation, with some exceptions (Figure 5b)

Although the two formulations vary in sample size, they have a similar average Silhouette score (around values of 0.25), which agrees with the UMAP visualization (partly overlapping but distinguishable groups). Moreover, the CV, MCV, and correlation values show a certain similarity between these populations in terms of ‘heterogeneity’ (of both single features and the full set of features) and in terms of feature correlations.

Similarly, when comparing formulations with medium-concentration of antibody functionalization and either 100 nm or 300 nm nanoparticle diameter (Figure 5e-h), the two formulations clearly separate with some overlap. This is visible from both the UMAP and the Silhouette scores. These scores have a similar average, with the 100 nm diameter formulation being more ‘compact’ in the feature space, overlapping more with the 300 nm diameter formulation and hence having lower Silhouette scores. While these two populations show similar low variability of their features (as visible from the MCV and CVs), they show different values of their feature correlations.

To prove the usefulness of these tools in a relevant scenario, we applied this approach to assess reproducibility, *e*.*g*., between batches of the same formulation. This is crucial in both the academic setting (to ensure the robustness of data published and validated by other groups) and the industrial setting (to assess batch-to-batch variations that may be detrimental to the quality of the product). Here we showcase how the metrics can be used to assess reproducibility between batches of the same formulation obtained: (a) by different operators, and (b) by the same operator, at differing formulation and analysis times.

When looking at different operators, here indicated as operator “green” and “purple” (Figure 6a-e), we focused on the formulation with medium-concentration of antibody functionalization and 200 nm nanoparticle diameter (200 medium), through multiple days. The UMAP plot (Figure 6c) shows that the samples of the two operators differ in heterogeneity, with the “purple” operator consistently producing more heterogeneous nanoparticles. This is also reflected in the silhouette scores (Figure 6b), with the “green” operator reaching Silhouette scores mostly between 0.25 and 0.50, while the “purple” operator has Silhouette scores ranging from 0.15 to *-* 0.50 (indicating a more ‘dispersed’ feature space).

**Figure 6:**
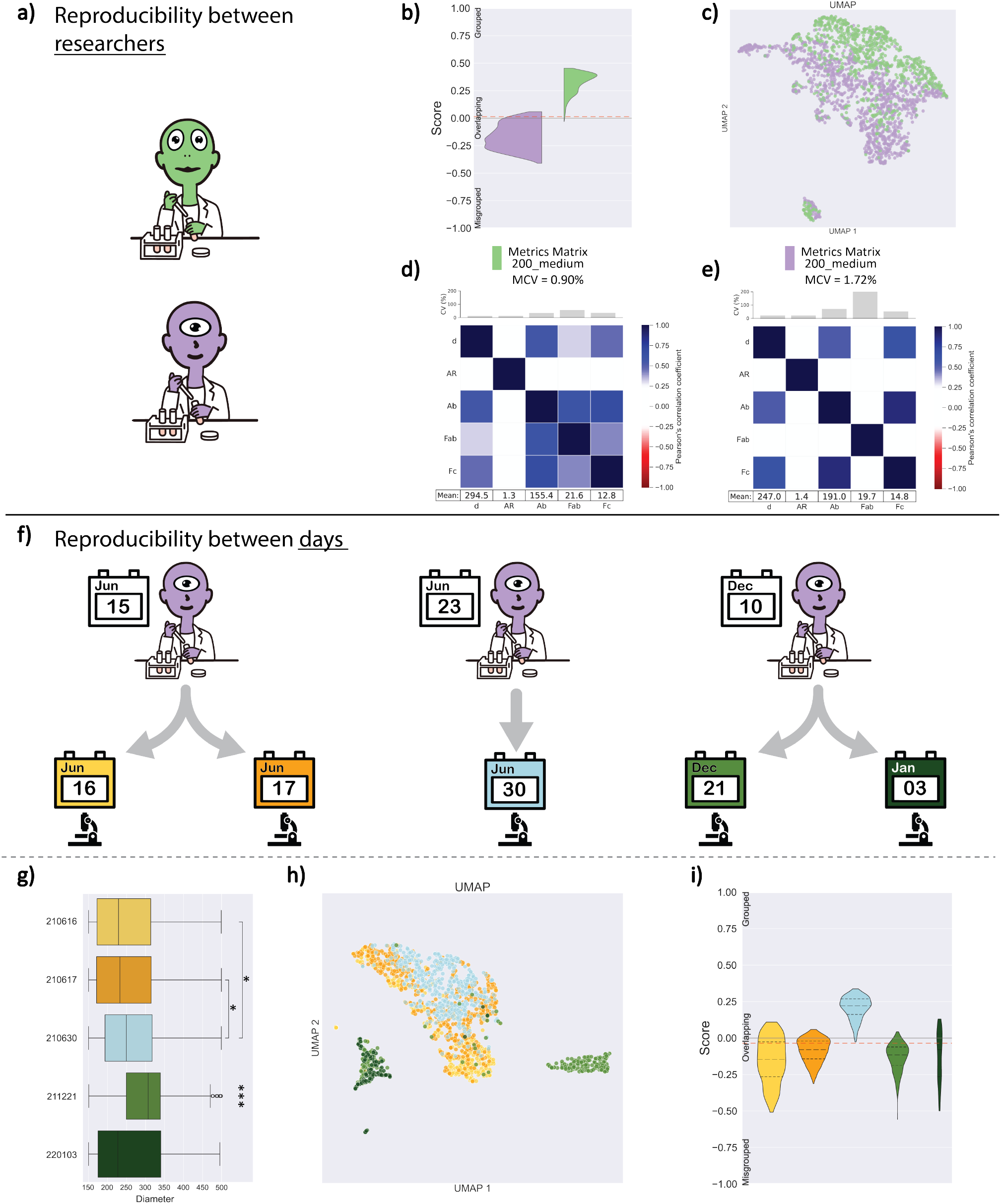
Reproducibility study of nanoparticle formulations. **a-e)** Between two different operators, and **f-i** by the same operator over time. **a)** Two operators synthesize and image the nanoparticle formulation for medium-concentration of antibody functionalization and 200 nm diameter (200 medium), at different times. **b)** Silhouette scores comparison of both reproductions for formulation 200 medium. **c)** UMAP plot calculated for both reproductions. **d-e)** Pearson’s correlation matrices, with the MCV per formulation and CV per formulation and feature. **f)** One operator synthesizes and images the nanoparticle formulation with a medium concentration of antibody functionalization and 300 nm diameter (300 medium), with multiple reproductions across seven months. All experimental reproductions of this formulation were synthesized starting from the same commercial nanoparticles batch. The first nanoparticle formulations were synthesized on June 15^th^, the second on June 23^rd^, and the third on December 10^th^. Then, the same operator imaged these multiple reproductions on different days: the first on June 16^th^ and 17^th^, the second on June 30^th^, and the third on December 21^st^ and January 3^rd^. **g)** Boxplot showing the nanoparticle diameter measured with super-resolution microscopy. Significant differences were found between some reproductions, marked with an * at *p <* 0.05 and *** at *p <* 0.0005. **h)** UMAP visualization of the multiple reproductions. **i)** Silhouette scores for each of the reproductions.

In any case, most of the nanoparticles produced by the operators overlap, denoting similarity and thus reproducibility of a few samples. This is also visible in similar trends in their CV and MCV values (Figure 6d and e). The differences in the nanoparticle dispersion across operators can be elucidated by both the CV and the correlation matrix, which can be used to pinpoint specific differences within features (CV) and across features (correlation). Finally, in Supplementary Figure S20, we can observe how the two replicas of the same formulation are more similar to each other (overlapping) than to a different formulation (grouped separately).

To study the reproducibility of the experiments over time, one operator was tasked with synthesizing one formulation (300 medium) at different time points, over a time span of seven months. These batches were then imaged at different intervals from the formulation date (ranging from one day to 24 days after formulation, (Figure 6f). In Figure 6g, we can observe the variance of the nanoparticle’s diameter, a common feature used to test the reproducibility of an experiment. However, when looking at one feature at a time (*e*.*g*., the diameter), we found a few formulations with statistically significant differences (Kruskal-Wallis H-test and Bonferroni test as a post hoc); formulation 211221 shows statistically significant differences with all other replicas with a p-value of 0.0005, and the pairs 210616-210630 and 210617-210630 with a p-value of 0.05. However, when observing the samples in a multivariate way, statistically significant differences also emerge in another replica (220103). When observing the UMAP (Figure 6h), batches synthesized later in the year, they grouped separately from the rest, with a few exceptions. The same trend is visible in the Silhouette scores (Figure 6i), where the data of the first batch, made from fresh material, has a similar score distribution and is shown to overlap. Then the nanoparticle batch made later in the same month follows a similar trend, although it clusters slightly separated but within the same region as the oranges. Finally, the batch made months later can be observed to have an overall lower silhouette score. Finally, the samples deteriorate over time (*e*.*g*., the batch prepared in December that was imaged in January). These aspects are also visible when observing all metrics introduced thus far (Supplementary Figure S19), noting how the MCV value (and thus the heterogeneity of the sample) increases over time.

These results show how the multivariate analysis of formulations, along with the metrics introduced in this work, can shed light on reproducibility, which would not be possible otherwise (*e*.*g*., by looking at one feature at a time). Indicating which features vary from batch to batch opens unprecedented opportunities to guide the optimization and standardization of synthetic procedures.

## 3 Conclusions and Outlook

In this work, we introduced a comprehensive nanoinformatics framework for the analysis of complex nanoparticle datasets. Our approach addresses several key challenges in the field of nanomaterial characterization by providing tools to visualize, describe, and compare nanoparticle formulations at the single-particle level. By applying diverse dimensionality reduction algorithms (PCA, t-SNE, UMAP, and MST) to multivariate super-resolution microscopy data, we have demonstrated how complex nanoparticle libraries can be effectively visualized while preserving crucial information about their heterogeneity and (dis)similarities. Moreover, we also introduce quantitative metrics to describe nanoparticle formulations. By adapting established statistical measures to the multivariate nature of nanomaterials data, we have developed a “nanoparticle ID card” that captures both individual feature variations (through the CV) and global formulation heterogeneity (through the MCV). This approach provides a more complete understanding of the nanoparticle characteristics in comparison to traditional single-parameter analyses, enabling researchers to comprehend the complex interactions among material properties. Furthermore, we have demonstrated how the Silhouette score can be used as a powerful tool to compare different nanoparticle formulations and assess reproducibility. Combined with our visualization and description framework, this metric provides unprecedented insight into batch-to-batch variations and operator-dependent differences in nanoparticle synthesis. Our analysis revealed how multivariate characterization can uncover subtle differences between formulations that might be missed by traditional single-parameter analyses, highlighting the importance of comprehensive characterization approaches in nanomaterial development. The tools presented here represent a step toward standardizing the analysis of complex nanomaterial data. By providing a systematic framework for characterizing and comparing nanoparticle formulations, our approach can facilitate the following:

- More robust quality control in nanomaterial production
- Better understanding of structure-property relationships
- More effective optimization of synthetic procedures
- Enhanced reproducibility across different laboratories
- More informed decision-making in nanomaterial design

Looking forward, this framework can be extended to other characterization techniques and material types, potentially serving as a foundation for a broader materials-omics approach. As data-driven approaches for nanomaterials continue to grow, standardized methods for analyzing and comparing complex datasets will become increasingly important. Our work provides a roadmap for how such analyses can be performed, potentially accelerating the development and application of nanomaterials across various fields. These advances in nanoinformatics, combined with multiparametric characterization methods such as super-resolution microscopy, are positioning the field to take advantage of modern machine learning and artificial intelligence tools, potentially leading to more rational and efficient approaches to materials design and optimization.

## 4 Materials and Methods

### 4.1 Dataset

The nanoparticle library composed of silica nanoparticle formulations, used in this work, was synthesized and characterized by PAINT super-resolution microscopy, as shown by Tholen et al. [29]. Then, the dataset was reanalyzed to obtain twenty-three features on a single-particle level using the *nanoFeatures* app [36].

### 4.2 Uni- and multivariate metrics

#### Pearson Correlation coefficient

Given two features, their pairwise Pearson’s correlation coefficient (*r*) is defined as follows:

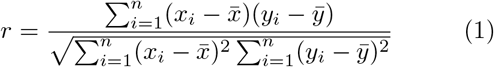

where *x*_*i*_ is the *i*-th data point of the first variable *X, y*_*i*_ is the *i*-th data point of the second variable *Y*, 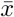 is the mean of the first variable *X*, 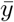 is the mean of the second variable *Y*, *n* is the total number of data points. *r* quantifies the strength and direction of the linear relationship between two variables, and it ranges from −1 (perfect negative linear relationships) to +1 (perfect positive linear relationship), with *r* = 0 indicating no linear correlation.

#### Polydispersity Index

The Polydispersity Index (PDI) for the *i*-th feature can be computed as follows:

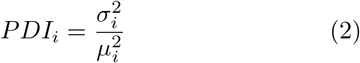

where *σ*_*i*_ is the standard deviation of the *i*-th particle size distribution, and *µ*_*i*_ is the corresponding mean. PDI ranges from 0 (monodisperse) to values greater than 0.4 (polydisperse), with 0.1 to 0.4 indicating moderate polydispersity.

#### Coefficient of Variation

The coefficient of variation (CV) for the *i*-th feature can be computed as follows:

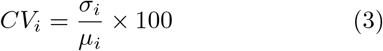

where *σ*_*i*_ is the standard deviation of the *i*-th feature, and *µ*_*i*_ is the mean of the *i*-th feature. The CV ranges from 0 to infinity; the higher the CV, the greater the relative variability of the feature compared to its mean.

#### Multivariate Coefficient of Variation

The Multivariate Coefficient of Variation (MCV) extends the CV to the multivariate scenario. In this work, the MCV was computed in agreement with [52], as follows:

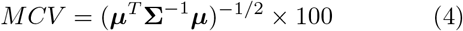

where ***µ*** is a vector of means for all features being considered, and **Σ** is the covariance matrix of the features (which captures the relationships and co-variability between the features). The MCV ranges from 0 to infinity; the higher the MCV, the greater the relative variability of the data in relation to the mean. A higher MCV indicates that the data is more ‘spread out’ in the variables space, while a lower MCV suggests a lower relative variability.

This MCV formulation cannot be calculated when the covariance matrix is singular (*i*.*e*., it cannot be inverted). In those (unlikely) cases, MCV is assigned a ‘dummy’ value of MCV = −1.

#### 4.2.1 Silhouette Scores

Given two nanoparticle formulations, – *a*, and *b* – the Silhouette scores [53] were computed as:

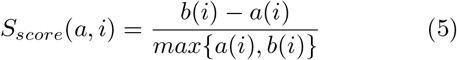

Where *i* is a nanoparticle belonging to the formulation *a*(*i*) is the average of the distances of *i* to all the other nanoparticles of the same formulation, and *b*(*i*) is the average of the Euclidean distances from nanoparticle *i* to all the other nanoparticles of the formulation In the case of multiple formulations, formulation *b* will be the closest to *a*, that is, the formulation with the minimum average distance to *a*. In other words, the Silhouette score compares (a) the similarity of the *i*-th nanoparticle to all nanoparticles of the same formulation (*a(i)*), with (b) its similarity to the other formulation (*b*(*i*)) – providing a measure of how well each nanoparticle matches the characteristics of the other particles of the same formulation. *S*_*score*_ ranges between −1 (nanoparticle is more similar to its closest formulation than its own), 0 (formulations perfectly overlap), and 1 (perfect separations).

In this work, the *S*_*score*_ were calculated directly on the dataset features, instead of on the clustered data,

### 4.3 Hyperparameter tuning

The hyperparameters for t-SNE were optimized to minimize the Kullback-Leibler divergence, as detailed in Table S2, which resulted in a Kullback-Leibler divergence of 0.997 and an effective learning rate of 435.875. For UMAP, the hyperparameters chosen are specified in Table S3, these were selected to optimize formulation separation and minimize computational time. Supplementary Figures S7 and S8 illustrate the effects of parameter selection.

### 4.4 Software and Code

Data analysis was performed using Python (v3.12.3) with SciPy (v1.13.0) and NumPy (v1.26.4). We used scikit-learn (v1.4.2) to apply the PCA (initialized with random state = 2023) and t-SNE. We used umap-learn (v0.5.6)[47] for UMAP.

Finally, in this work, we have used the MST implementation in SciPy v1.13.0, which uses Kruskal’s algorithm [49] to calculate the tree. The algorithm connects every single nanoparticle from the different samples to its closest neighbor, based on the Euclidean distance between their features.

## 5 Author contributions

*Conceptualization*: CIL, LA, and FG. *Resources:* AS and MM. *Data curation:* CIL, MMET, and VG. *Formal analysis:* CIL, MMET, and VG. *Methodology:* CIL, FG, and LA. *Software:* CIL. *Visualization:* CIL and MMET. *Writing - original draft:* CIL and MMET. *Writing - review and editing:* all authors. All authors have approved the final version of the manuscript.

## 7 Code availability

The code and data used in this work can be found in GitHub at the following URL: https://github.com/n4nlab/npAtlas. Moreover, the repository contains an interactive version of the UMAP plot to facilitate data analysis and interpretation, available as a Jupyter Notebook in the aforementioned GitHub repository.

## 8 Acknowledgements

Schematic illustrations were created using Biorender.com. MMET and LA appreciate the financial support from The Netherlands Organization for Scientific Research (NWO VIDI Grant 192.028).

## References

[1] Edgar López-López, Jurgen Bajorath, and José L Medina-Franco. “Informatics for chemistry, biology, and biomedical sciences”. In: Journal of chemical information and modeling 61.1 (2020), pp. 26–35.

[2] Arsalan Wafi and Reza Mirnezami. “Translational–omics: future potential and current challenges in precision medicine”. In: Methods 151 (2018), pp. 3–11.

[3] Syed Abdul Majeed Musavi et al. “Omics Technology: Revolution in Plant Biology”. In: Principles and Practices of OMICS and Genome Editing for Crop Improvement. Springer, 2022, pp. 197–212.

[4] Mathias Uhlén et al. “Tissue-based map of the human proteome”. In: Science 347.6220 (2015), p. 1260419.

[5] Tracy Hampton. “Cancer genome atlas”. In: Jama 296.16 (2006), pp. 1958–1958.

[6] The Cancer Genome Atlas Program (TCGA) - NCI. Archive Location: nciglobal,ncienterprise. May 13, 2022. url: http://www.cancer.gov/ccg/research/genome-sequencing/tcga.

[7] Andreas Digre and Cecilia Lindskog. “The human protein atlas—Integrated omics for single cell mapping of the human proteome”. In: Protein Science 32.2 (2023), e4562.

[8] Jeremy T-H Chang, Yee Ming Lee, and R Stephanie Huang. “The impact of the Cancer Genome Atlas on lung cancer”. In: Translational Research 166.6 (2015), pp. 568–585.

[9] Joerg Kurt Wegner et al. “Cheminformatics”. In: Communications of the ACM 55.11 (2012), pp. 65–75.

[10] David S Wishart. “Introduction to cheminformatics”. In: Current protocols in bioinformatics 53.1 (2016), pp. 14–1.

[11] Hongming Chen, Thierry Kogej, and Ola Engkvist. “Cheminformatics in drug discovery, an industrial perspective”. In: Molecular Informatics 37.9-10 (2018), p. 1800041.

[12] Xi Chen, Ming Liu, and Michael K Gilson. “BindingDB: a web-accessible molecular recognition database”. In: Combinatorial chemistry & high throughput screening 4.8 (2001), pp. 719–725.

[13] Michael K Gilson et al. “BindingDB in 2015: a public database for medicinal chemistry, computational chemistry and systems pharmacology”. In: Nucleic acids research 44.D1 (2016), pp. D1045– D1053.

[14] Anne Mai Wassermann and Jürgen Bajorath. “BindingDB and ChEMBL: online compound databases for drug discovery”. In: Expert opinion on drug discovery 6.7 (2011), pp. 683–687.

[15] Barbara Zdrazil et al. “The ChEMBL Database in 2023: a drug discovery platform spanning multiple bioactivity data types and time periods”. In: Nucleic acids research 52.D1 (2024), pp. D1180– D1192.

[16] Liya Feng et al. “Small molecule drug discovery for glioblastoma treatment based on bioinformatics and cheminformatics approaches”. In: Frontiers in Pharmacology 15 (2024), p. 1389440.

[17] Ewelina Wyrzykowska et al. “Representing and describing nanomaterials in predictive nanoinformatics”. In: Nature Nanotechnology 17.9 (2022), pp. 924–932.

[18] Kurt LM Drew et al. “Size estimation of chemical space: how big is it?” In: Journal of Pharmacy and Pharmacology 64.4 (2012), pp. 490–495.

[19] Ghanshyam Pilania et al. “Accelerating materials property predictions using machine learning”. In: Scientific reports 3.1 (2013), p. 2810.

[20] Yongtae Kim et al. “Deep learning framework for material design space exploration using active transfer learning and data augmentation”. In: npj Computational Materials 7.1 (2021), p. 140.

[21] Mahmoud Nasrollahzadeh et al. “Types of nanostructures”. In: Interface science and technology 28 (2019), pp. 29–80.

[22] Atsuto Seko et al. “Progress in nanoinformatics and informational materials science”. In: MRS Bulletin 43.9 (2018), pp. 690–695.

[23] Fernando González-Nilo et al. “Nanoinformatics: an emerging area of information technology at the intersection of bioinformatics, computational chemistry and nanobiotechnology”. In: Biological Research 44.1 (2011), pp. 43–51.

[24] Teodora Andrian et al. “Correlating super-resolution microscopy and transmission electron microscopy reveals multiparametric heterogeneity in nanoparticles”. In: Nano Letters 21.12 (2021), pp. 5360–5368.

[25] Yuyang Wang et al. “Multicolor super-resolution microscopy of protein corona on single nanoparticles”. In: ACS Applied Materials & Interfaces 14.33 (2022), pp. 37345–37355.

[26] Laura Woythe et al. “A single-molecule view at nanoparticle targeting selectivity: Correlating ligand functionality and cell receptor density”. In: ACS nano 16.3 (2022), pp. 3785–3796.

[27] Laura Woythe et al. “Single-particle functionality imaging of antibody-conjugated nanoparticles in complex media”. In: ACS Applied Bio Materials 6.1 (2023), pp. 171–181.

[28] Pietro Delcanale et al. “Nanoscale mapping functional sites on nanoparticles by points accumulation for imaging in nanoscale topography (PAINT)”. In: ACS nano 12.8 (2018), pp. 7629–7637.

[29] Marrit ME Tholen et al. “Mapping antibody domain exposure on nanoparticle surfaces using dna-paint”. In: ACS nano 17.12 (2023), pp. 11665–11678.

[30] Emmanouil Archontakis et al. “Spectrally PAINTing a single chain polymeric nanoparticle at super-resolution”. In: Journal of the American Chemical Society 144.51 (2022), pp. 23698–23707.

[31] Emmanouil Archontakis et al. “Mapping the relationship between total and functional antibodies conjugated to nanoparticles with spectrally-resolved direct stochastic optical reconstruction microscopy (SR-dSTORM)”. In: Nanoscale Advances 4.20 (2022), pp. 4402–4409.

[32] Teodora Andrian, Silvia Pujals, and Lorenzo Albertazzi. “Quantifying the effect of PEG architecture on nanoparticle ligand availability using DNA-PAINT”. In: Nanoscale Advances 3.24 (2021), pp. 6876–6881.

[33] Sara Cunha et al. “Using the quality by design (QbD) approach to optimize formulations of lipid nanoparticles and nanoemulsions: A review”. In: Nanomedicine: Nanotechnology, Biology and Medicine 28 (2020), p. 102206.

[34] Sergey K Filippov et al. “Dynamic light scattering and transmission electron microscopy in drug delivery: a roadmap for correct characterization of nanoparticles and interpretation of results”. In: Materials Horizons 10.12 (2023), pp. 5354–5370.

[35] Ralf Jungmann et al. “Quantitative super-resolution imaging with qPAINT”. In: Nature methods 13.5 (2016), pp. 439–442.

[36] Cristina Izquierdo Lozano et al. “nanoFeatures: a cross-platform application to characterize nanoparticles from super-resolution microscopy images”. In: Nanoscale (2024).

[37] Fleur Mougin et al. “Visualizing omics and clinical data: Which challenges for dealing with their variety?” In: Methods 132 (2018), pp. 3–18.

[38] Federico Taverna et al. “BIOMEX: an interactive workflow for (single cell) omics data interpretation and visualization”. In: Nucleic acids research 48.W1 (2020), W385–W394.

[39] Nils Gehlenborg et al. “Visualization of omics data for systems biology”. In: Nature methods 7.Suppl 3 (2010), S56–S68.

[40] Malte D Luecken and Fabian J Theis. “Current best practices in single-cell RNA-seq analysis: a tutorial”. In: Molecular systems biology 15.6 (2019), e8746.

[41] Yoon Ha Choi and Jong Kyoung Kim. “Dissecting cellular heterogeneity using single-cell RNA sequencing”. In: Molecules and cells 42.3 (2019), pp. 189–199.

[42] Dmitry Kobak and Philipp Berens. “The art of using t-SNE for single-cell transcriptomics”. In: Nature communications 10.1 (2019), p. 5416.

[43] Karl Pearson. “LIII. On lines and planes of closest fit to systems of points in space”. In: The London, Edinburgh, and Dublin philosophical magazine and journal of science 2.11 (1901), pp. 559–572.

[44] Svante Wold, Kim Esbensen, and Paul Geladi. “Principal component analysis”. In: Chemometrics and intelligent laboratory systems 2.1-3 (1987), pp. 37–52.

[45] Michael Greenacre et al. “Principal component analysis”. In: Nature Reviews Methods Primers 2.1 (2022), p. 100.

[46] Laurens Van der Maaten and Geoffrey Hinton. “Visualizing data using t-SNE.” In: Journal of machine learning research 9.11 (2008).

[47] Leland McInnes, John Healy, and James Melville. UMAP: Uniform Manifold Approximation and Projection for Dimension Reduction. 2020. url: https://arxiv.org/abs/1802.03426.

[48] Ali Grami. “Chapter 19 - Trees”. In: Discrete Mathematics. Ed. by Ali Grami. Academic Press, 2023, pp. 351–372. isbn: 978-0-12-820656-0. doi: 10.1016/B978-0-12-820656-0.00019-8. url: https://www.sciencedirect.com/science/article/pii/B9780128206560000198.

[49] Joseph B Kruskal. “On the shortest spanning subtree of a graph and the traveling salesman problem”. In: Proceedings of the American Mathematical society 7.1 (1956), pp. 48–50.

[50] Uwe Kruger, Junping Zhang, and Lei Xie. “Developments and applications of nonlinear principal component analysis–a review”. In: Principal manifolds for data visualization and dimension reduction (2008), pp. 1–43.

[51] Felix MD Marsh-Wakefield et al. “Making the most of high-dimensional cytometry data”. In: Immunology and Cell Biology 99.7 (2021), pp. 680–696.

[52] VG Voinov and MS Nikulin. Unbiased Estimators and their Applications Volume 2: Multivariate Case. 1997.

[53] Peter J Rousseeuw. “Silhouettes: a graphical aid to the interpretation and validation of cluster analysis”. In: Journal of computational and applied mathematics 20 (1987), pp. 53–65.

