## Supplementary material for "“*Visualize, describe, compare*” – nanoinformatics approaches for material-omics": Suplementary Information

*Table S1:* Nanoparticle library formulation list and description.

| Formulation name | Diameter (nm) | Amount of Ctx (Ctx/COOH) | Orientation | Color code |
|---|---|---|---|---|
| 100_medium | 100 | 0.68 | Random | |
| 100_high | 100 | 1.02 | Random | |
| 200_low | 200 | 0.34 | Random | |
| 200_medium | 200 | 0.68 | Random | |
| 200_high | 200 | 1.02 | Random | |
| 300_low | 300 | 0.34 | Random | |
| 300_medium | 300 | 0.68 | Random | |
| 300_high | 300 | 1.02 | Random | |
| 500_low | 500 | 0.34 | Random | |
| 500_medium | 500 | 0.68 | Random | |
| 500_high | 500 | 1.02 | Random | |
| NH2_pG0pM1 | 200 | 0.68 | 100% Fc | |
| NH2_pG1pM3 | 200 | 0.68 | 25% Fab, 75% Fc | |
| NH2_pG1pM1 | 200 | 0.68 | 50% Fab, 50% Fc | |
| NH2_pG3pM1 | 200 | 0.68 | 75% Fab, 25% Fc | |
| NH2_pG1pM0 | 200 | 0.68 | 100% Fab | |
| 200_medium_RPurple | 200 | 0.68 | Random | |
| 200_medium_RGreen | 200 | 0.68 | Random | |
| 300_medium_RPurple | 300 | 0.68 | Random | |
| 300_medium_RGreen | 300 | 0.68 | Random | |
| 300_medium_210616 | 300 | 0.68 | Random | |
| 300_medium_210617 | 300 | 0.68 | Random | |
| 300_medium_210630 | 300 | 0.68 | Random | |
| 300_medium_211221 | 300 | 0.68 | Random | |
| 300_medium_220103 | 300 | 0.68 | Random | |

#### 1 Dataset filtering

The data was filtered according to the total number of cluster localizations, and the quality of the fitting function to obtain the target counts, which is based on the probe’s binding kinetics. In the next figures, we compare the cluster target count (number of antibodies in the nanoparticles) found in each channel (Fab in channel 1 or Fc in channel 2) against the total number of localizations per cluster found in the image. These plots are repeated for each step in the dataset pre-processing. The dotted line indicates the diagonal, intuitively, (target count channel 1 + target count channel 2) ¡ total cluster localizations, since these are calculated by fitting a model on the cluster localizations. However, theoretically, in some special cases, these can be higher, for example in the case of high imager concentration. Therefore, filtering by the filtering quality should be the fairest for every dataset.

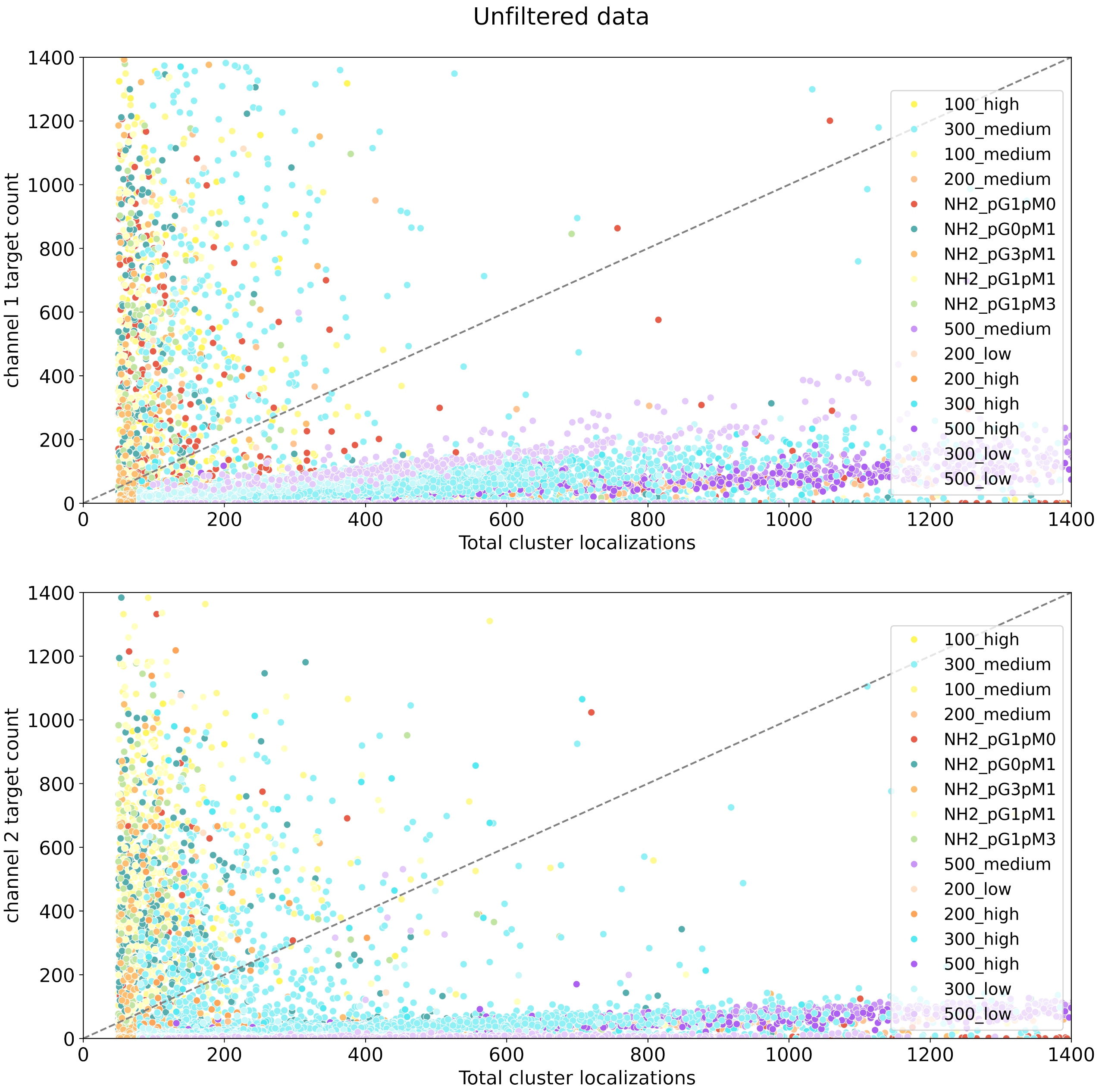


*Figure S1:* Comparison of the cluster target count per channel against the total number of localizations per cluster found in the image, for the unfiltered dataset.

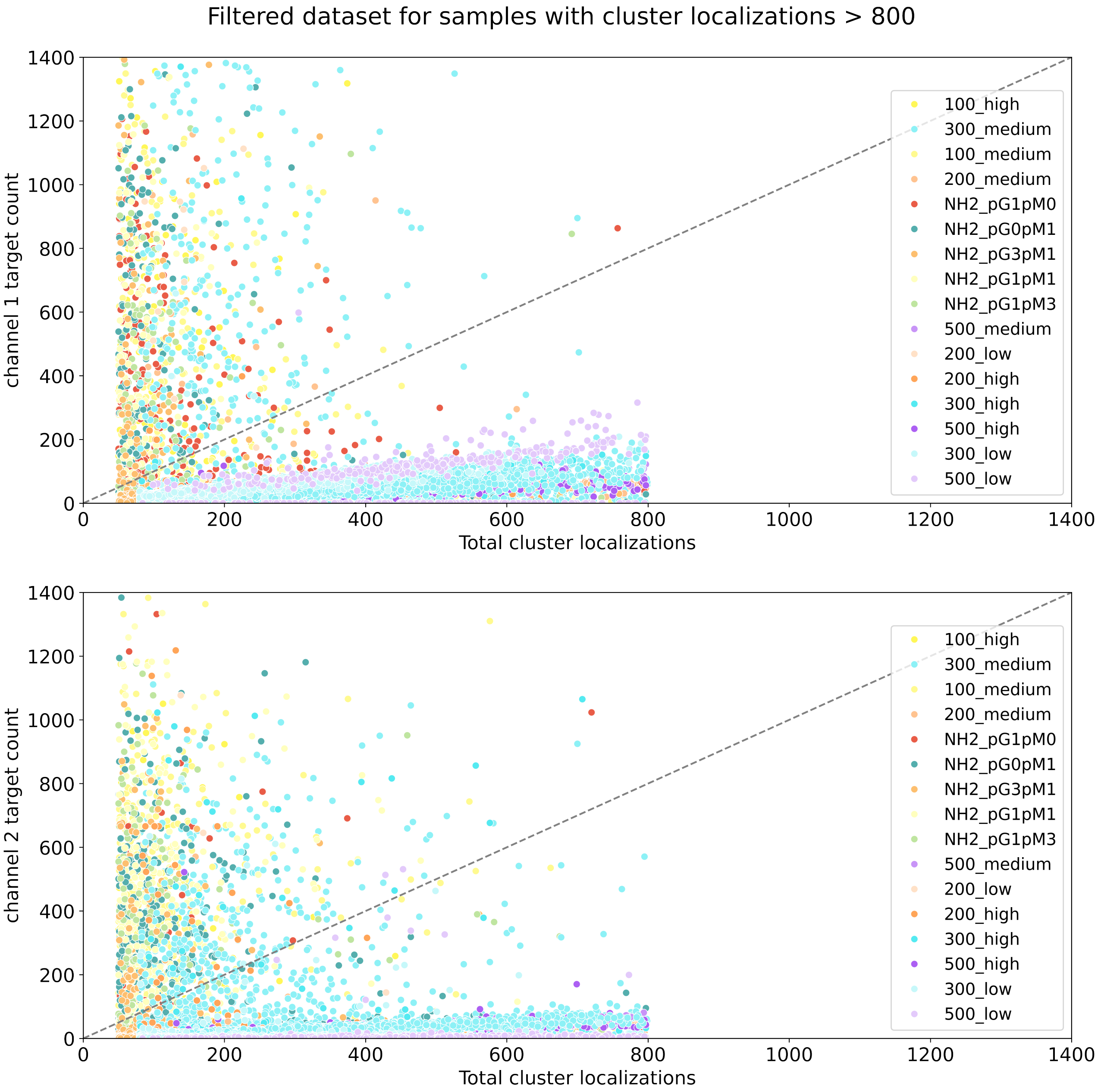


*Figure S2:* Comparison of the cluster target count per channel against the total number of localizations per cluster found in the image, for the dataset filtered by the number of total localizations.

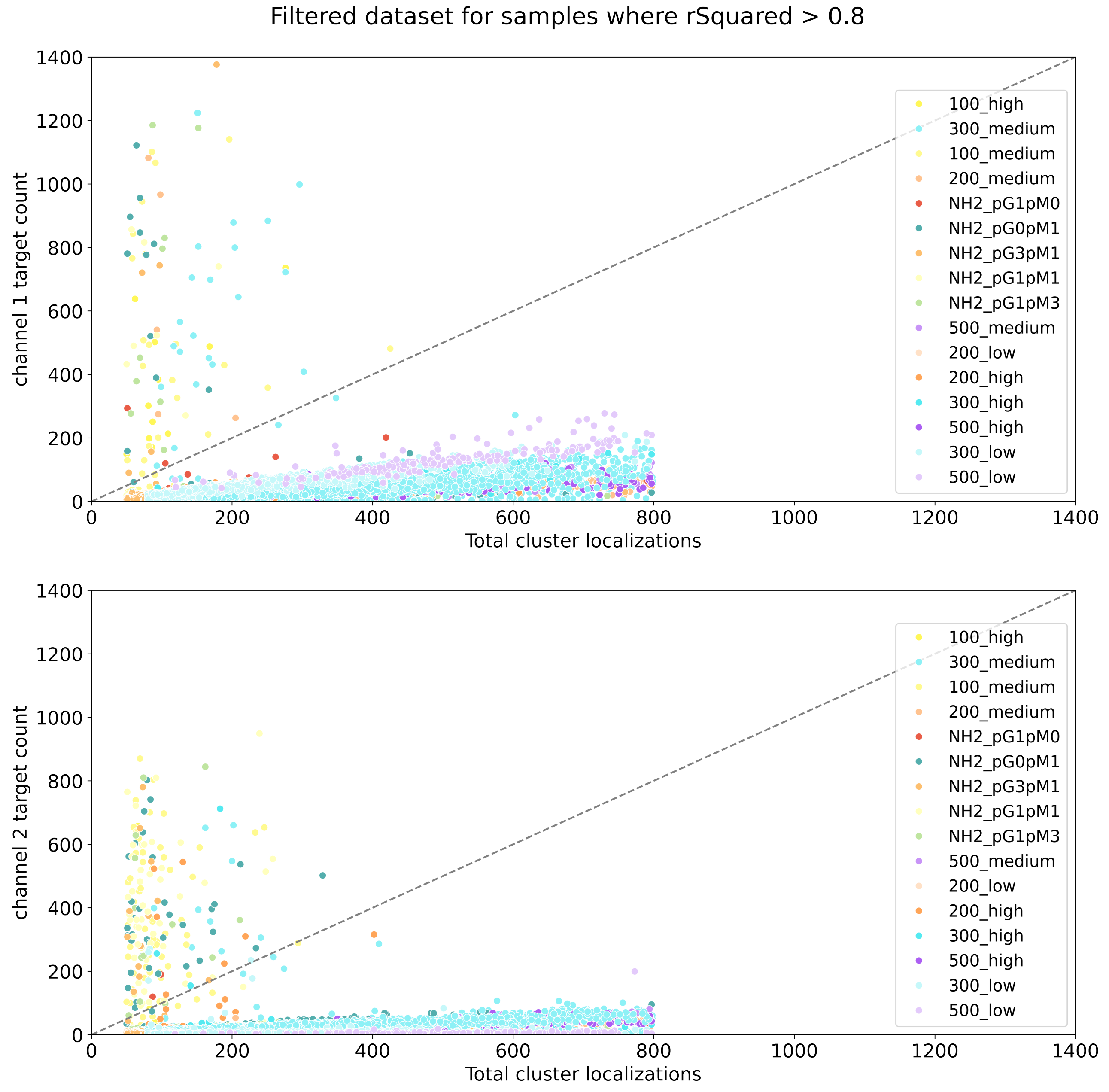


*Figure S3:* Comparison of the cluster target count per channel against the total number of localizations per cluster found in the image, for the dataset filtered by the number of total localizations and the fitting quality.

### 2 Metrics Matrix

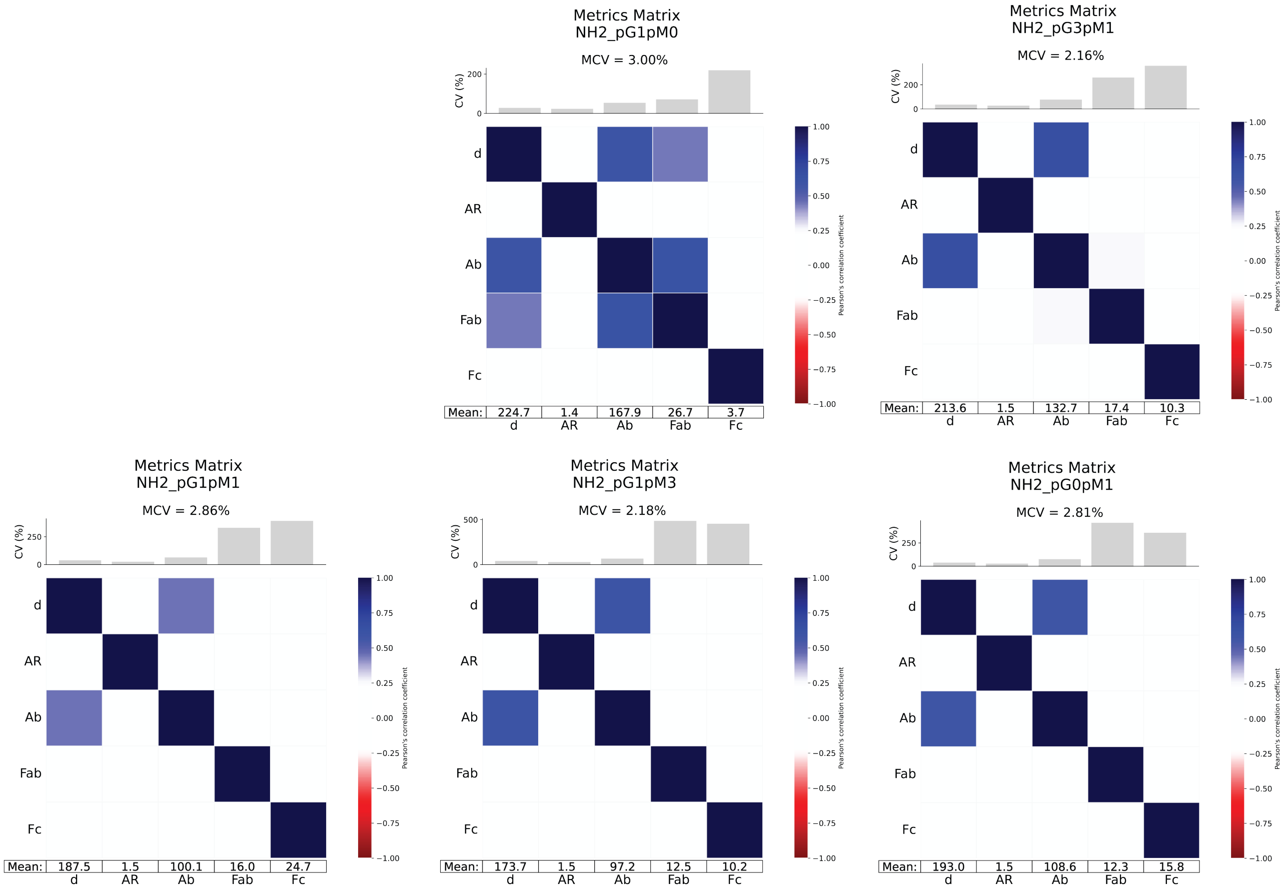


*Figure S4:* Metrics matrices for all formulations in the controlled antibody orientation NP formulations: varying ratio of protein type.

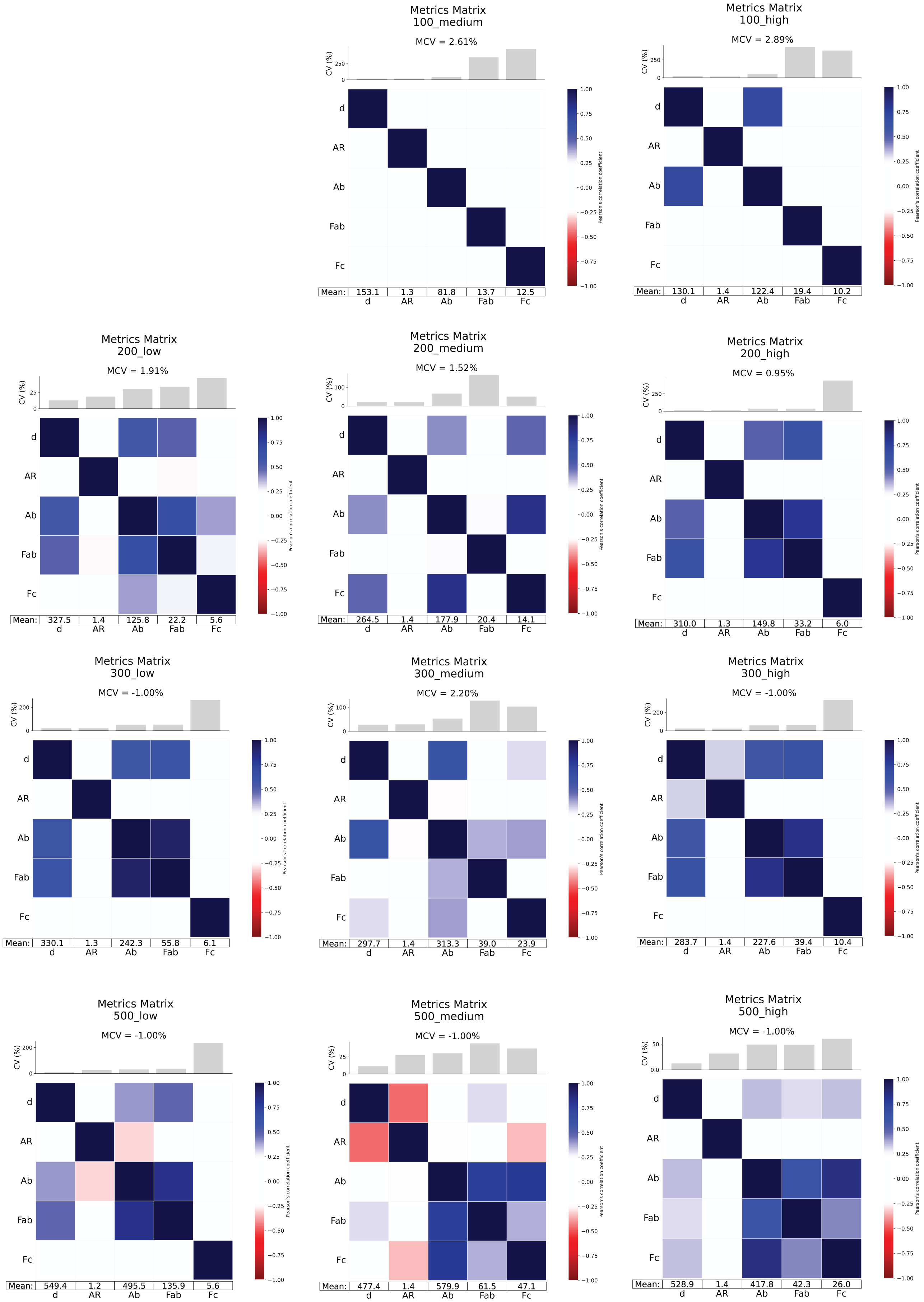


*Figure S5:* Metrics matrices for all formulations in the random antibody orientation NP formulations: varying size, and antibody on the surface concentration.

### 3 Dimensionality reduction algorithms

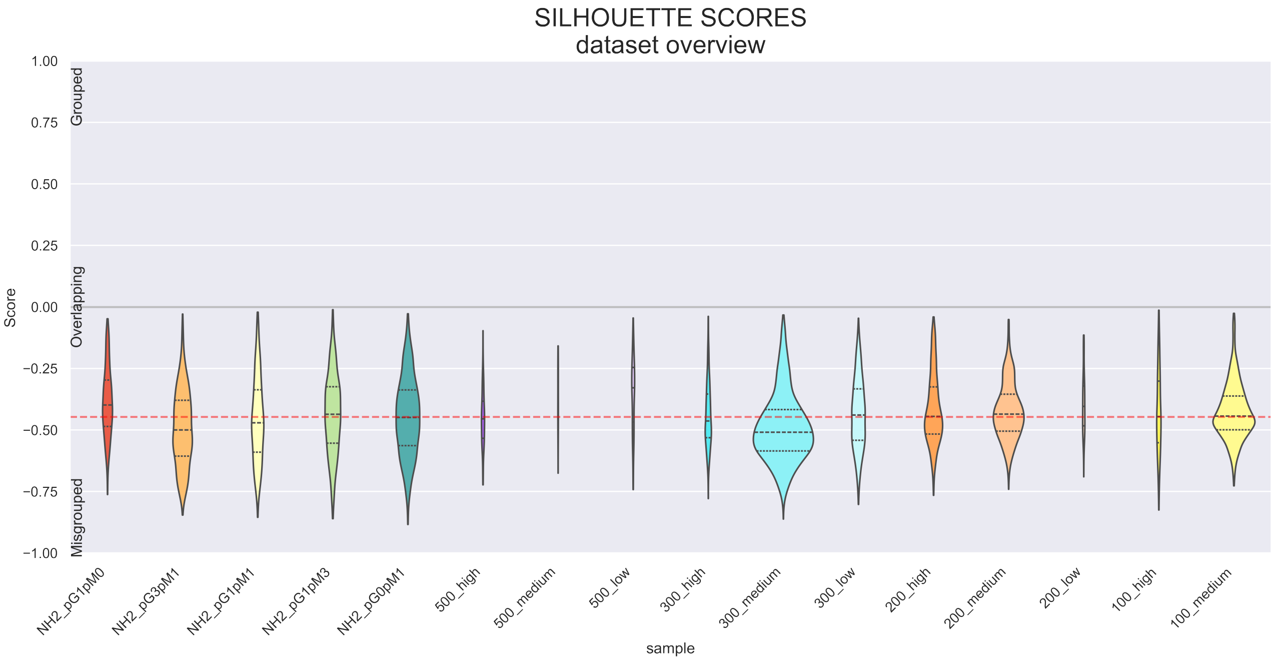


*Figure S6:* Silhouette scores of the full dataset, for each formulation. This score quantifies the clustering/grouping quality, meaning a value between -1 and 1 is calculated for each point. If it's -1 the point is misclustered/misgrouped, 0 overlaps with the closest cluster or formulation, and 1 is well clustered/grouped.

Both UMAP and t-SNE have many parameters that one can change in their Scikit-learn implementations. However, mainly the perplexity (for t-SNE) and the number of minimum neighbors and minimum distance (for UMAP) are used, since these hyperparameters affect the visualization the most. In the case of t-SNE, we can make use of the Kullback-Leibler (kl) divergence as a metric to choose the right perplexity value: the kl divergence should be minimized. Unfortunately, there is no such metric for UMAP, one needs to rely on visualizations and prior knowledge. In this case, we know that low values of minimum neighbors will focus on the local structure of the clusters (similarities within clusters), while higher values will focus the visualization on the global structure (similarities between clusters). In any case, different combinations of hyperparameters should be tested to properly interpret the graphs, as shown in the Supplementary Figures S7 and S8.

*Table S2:* Hyperparameters given for the chosen t-SNE visualization.

| **Hyperparameter** | **Value** |
|---|---|
| angle | 0.5 |
| early_exaggeration | 12.0 |
| init | pca |
| learning_rate | auto |
| method | barnes_hut |
| metric | euclidean |
| metric_params | None |
| min_grad_norm | 1e-07 |
| n_components | 2 |
| n_iter | 1000 |
| n_iter_without_progress | 300 |
| n_jobs | None |
| perplexity | 1000 |
| random_state | 42 |
| verbose | 1 |

*Table S3:* Hyperparameters given for the chosen UMAP visualization.

| **Hyperparameter** | **Value** |
|---|---|
| n_neighbors | 15 |
| n_epochs | 500 |
| init | spectral |
| learning_rate | 1 |
| min_dist | 0.1 |
| metric | euclidean |
| metric_kwds | None |
| output_metric | euclidean |
| output_metric_kwds | None |
| n_components | 2 |
| spread | 1 |
| low_memory | True |
| local_connectivity | 1 |
| repulsion_strength | 1 |
| negative_sample_rate | 5 |
| transform_queue_size | 4 |
| n_jobs | 1 |
| set_op_mix_ratio | 1 |
| angular_rp_forest | False |
| target_n_neighbors | -1 |
| target_metric | categorical |
| target_metric_kwds | None |
| target_weight | 0.5 |
| transform_seed | 42 |
| transform_mode | embedding |
| force_approximation_algorithm | False |

| | |
|---|---|
| unique | False |
| densmap | False |
| dens_lambda | 2 |
| dens_frac | 0.3 |
| dens_var_shift | 0.1 |
| output_dens | False |
| disconnection_distance | None |
| precomputed_knn | (None, None, None) |
| random_state | 25 |
| verbose | True |

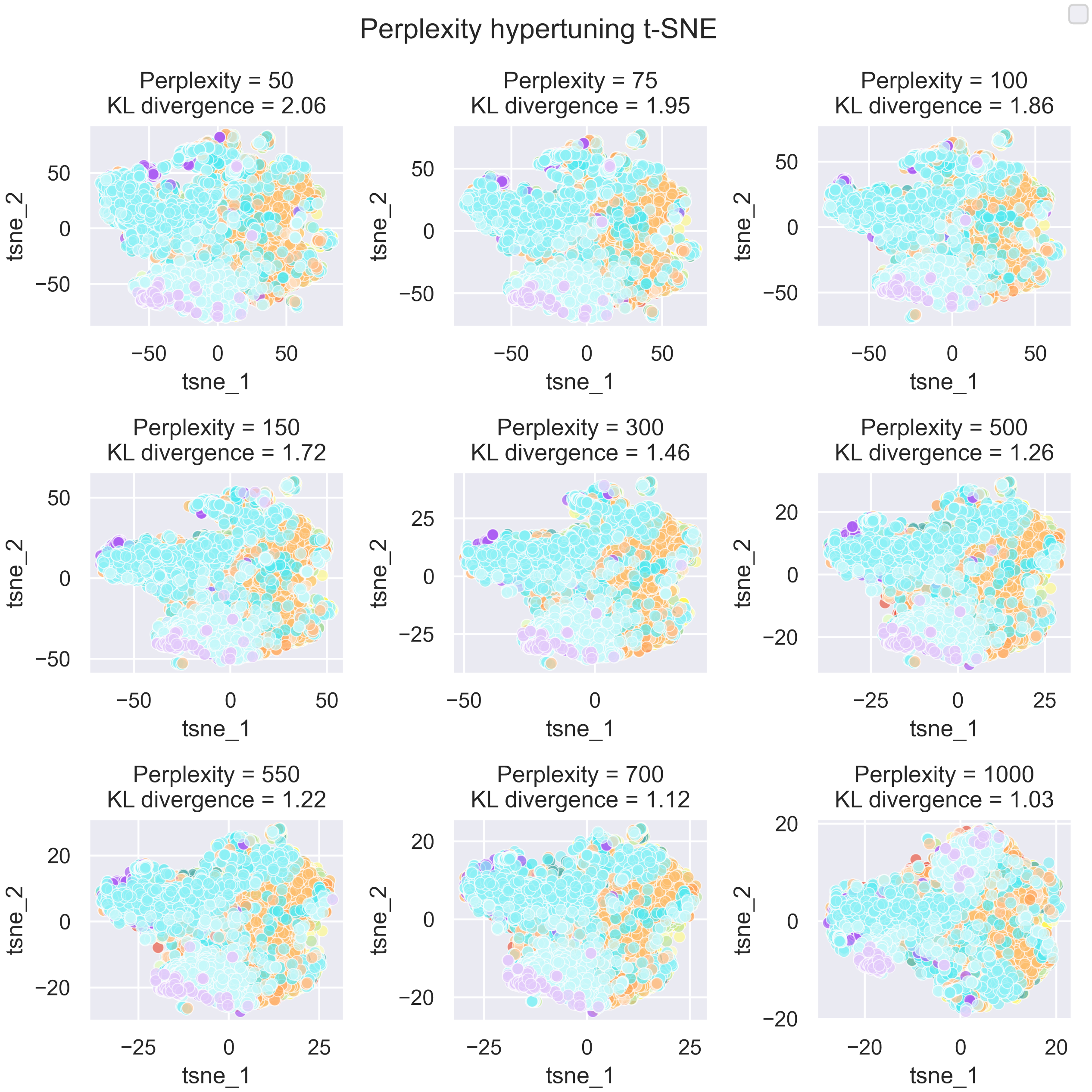


*Figure S7:* Hypertuning of the t-SNE perplexity value.

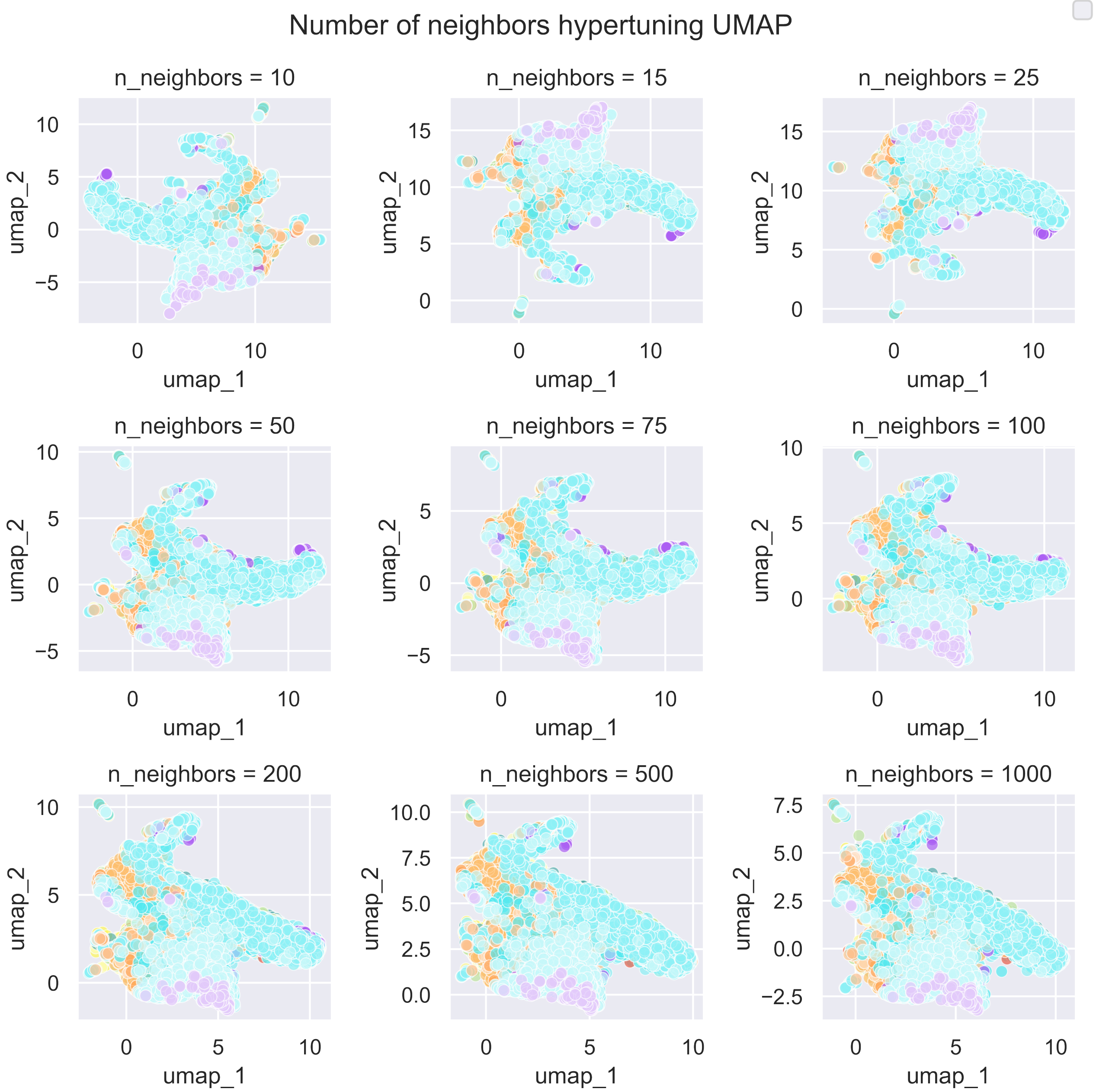


*Figure S8:* Hypertuning of the UMAP number of neighbors value.

The visualizations hardly vary when testing different tuning parameters, which may indicate a high similarity between populations. The chosen perplexity value for the t-SNE shown in the main text is 1000 as that value minimized the kl divergence, which is also the number of close neighbors expected for each cluster, since each formulation in the dataset contains approximately between 100 and 4000 samples. The number of neighbors and minimum distance chosen in the main text's UMAP are 15 and 0.1, respectively. The visualization was barely affected by these parameters, therefore, they were chosen to minimize computational time.

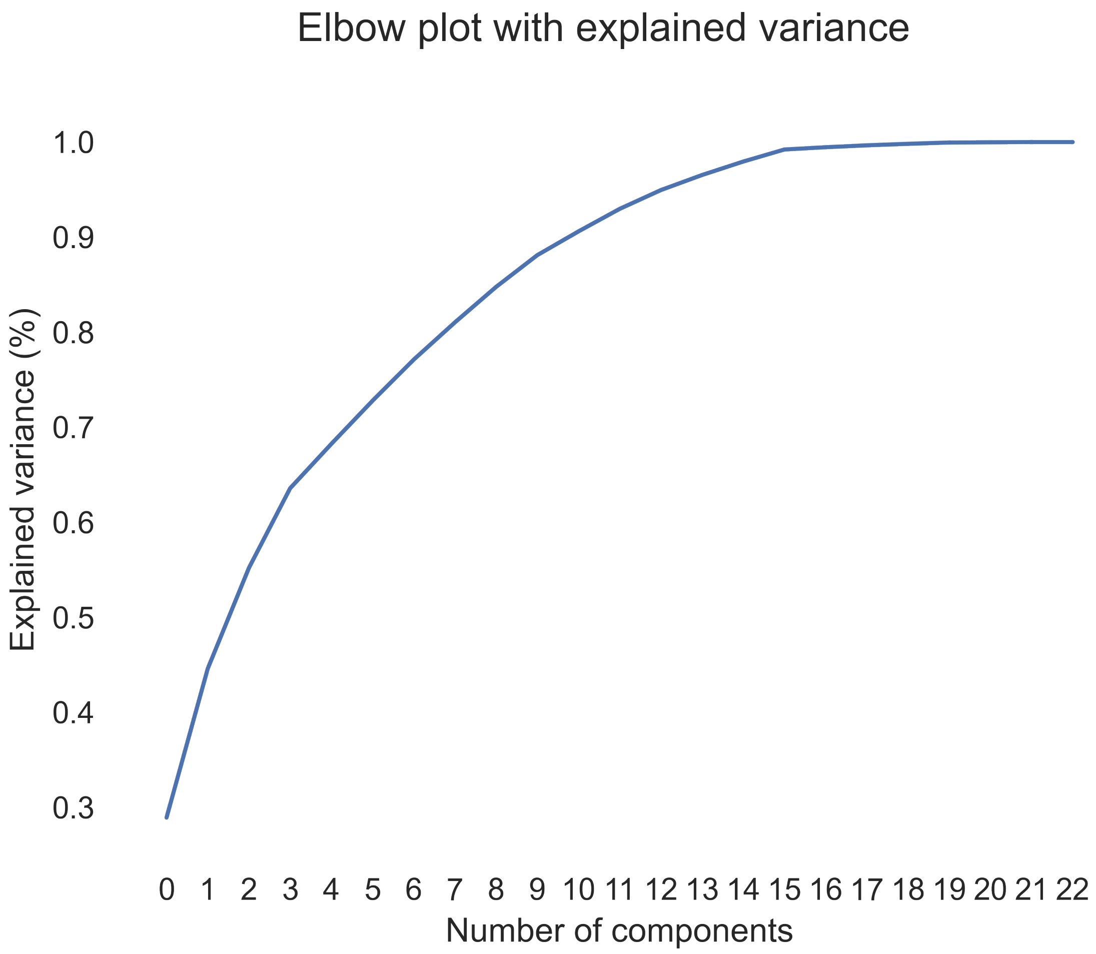


*Figure S9:* Elbow plot indicating the explained variance achieved by the principal components in the PCA.

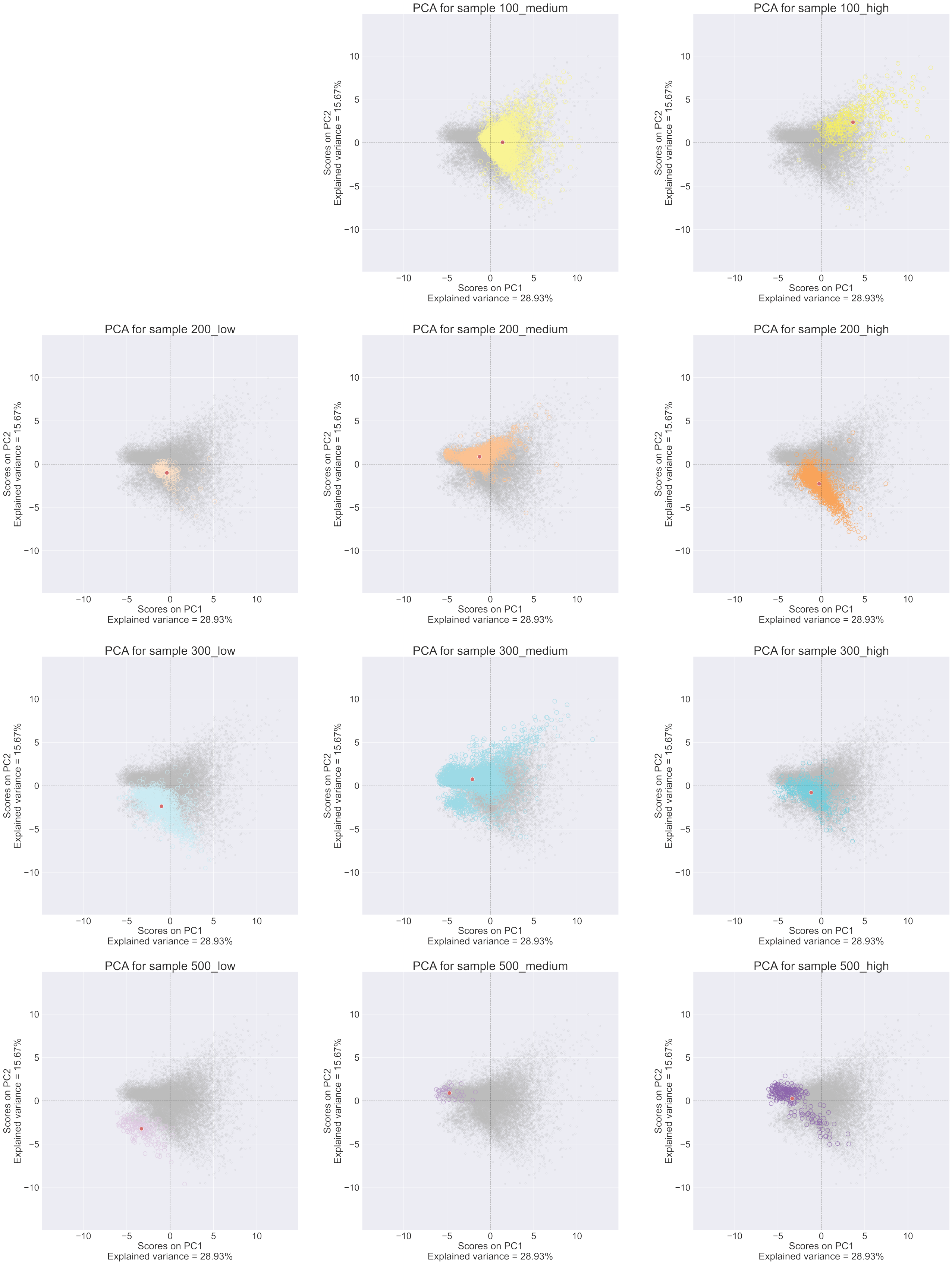


*Figure S10:* PCA plots where the color is highlighted for each formulation in the random antibody orientation NP formulations.

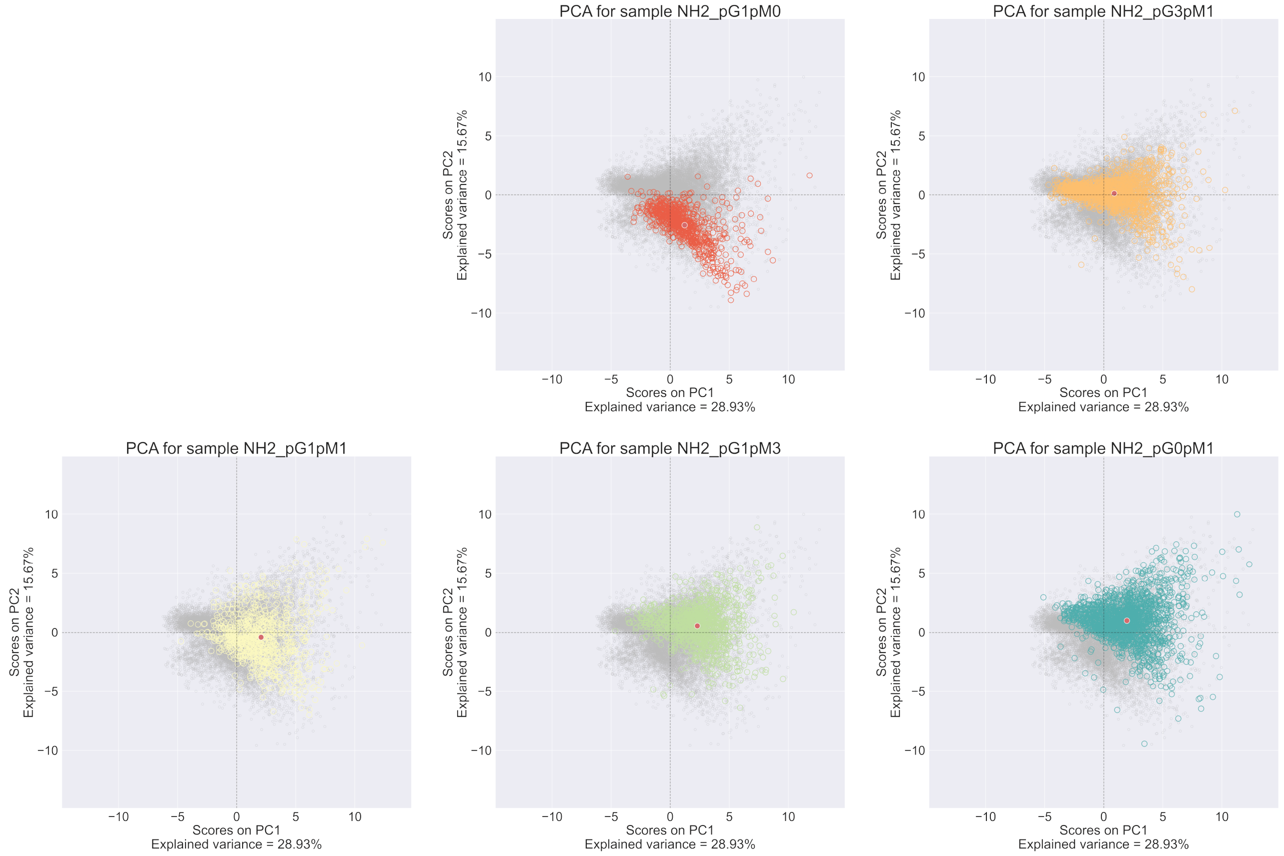


*Figure S11:* PCA plots where the color is highlighted for each formulation in the controlled antibody orientation NP formulations.

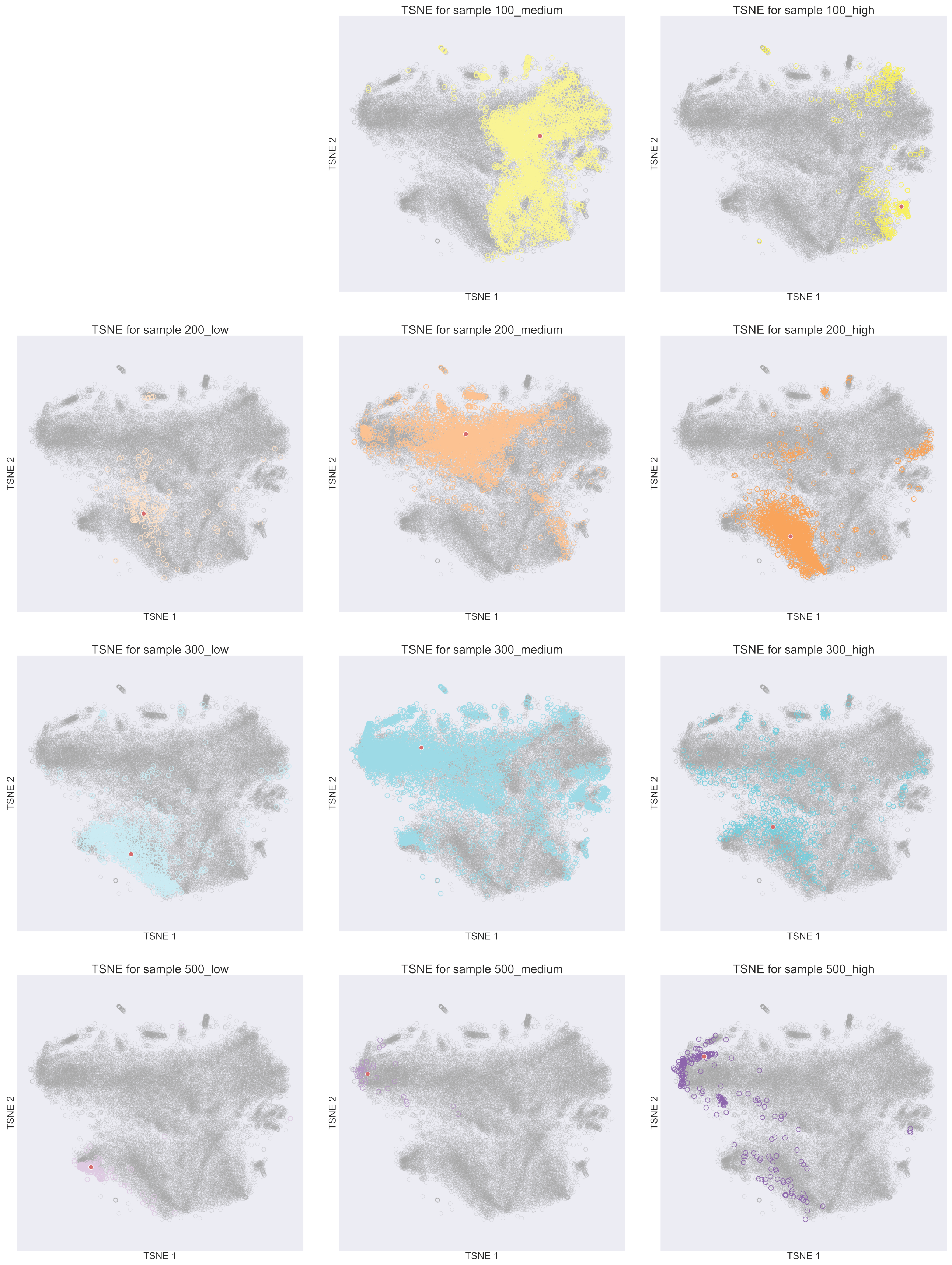


*Figure S12:* t-SNE plots where the color is highlighted for each formulation in the random antibody orientation NP formulations.

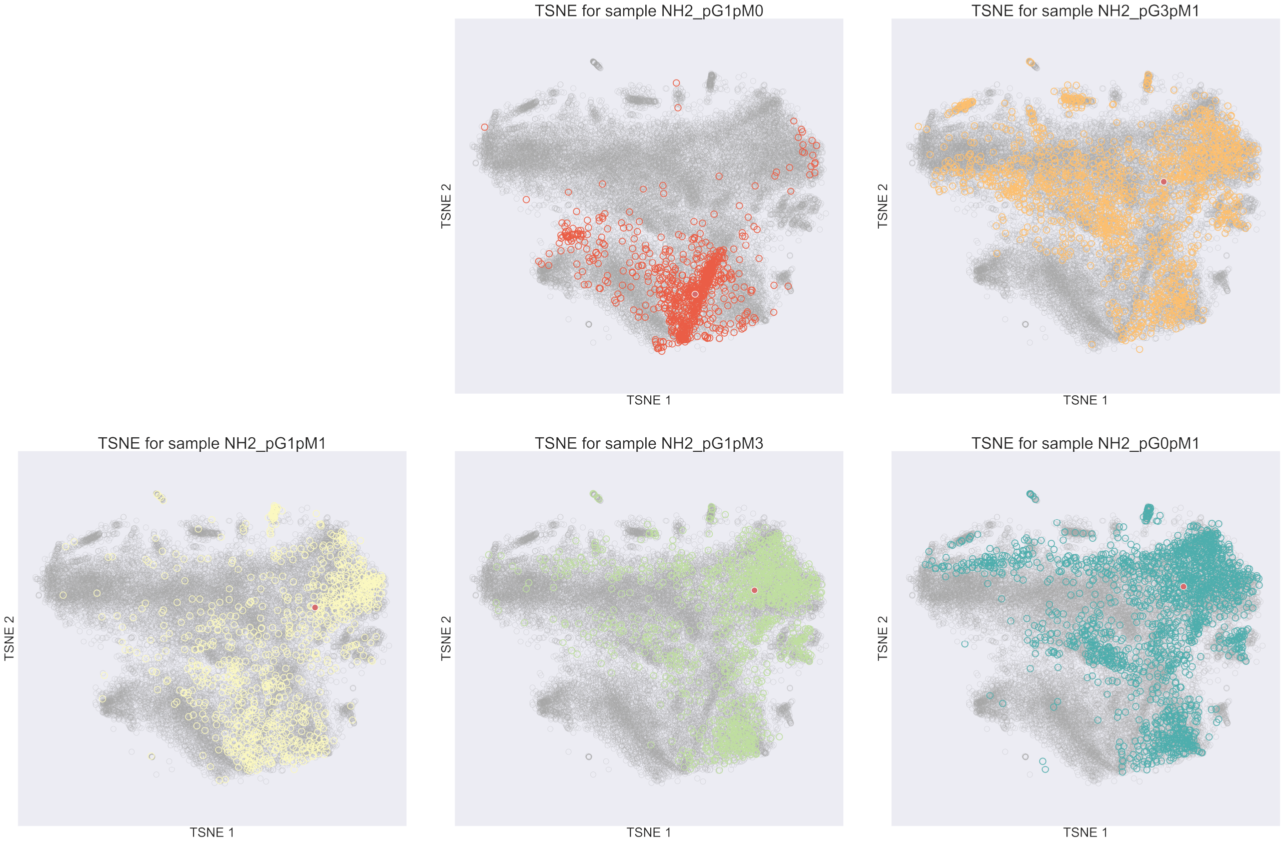


*Figure S13:* t-SNE plots where the color is highlighted for each formulation in the controlled antibody orientation NP formulations.

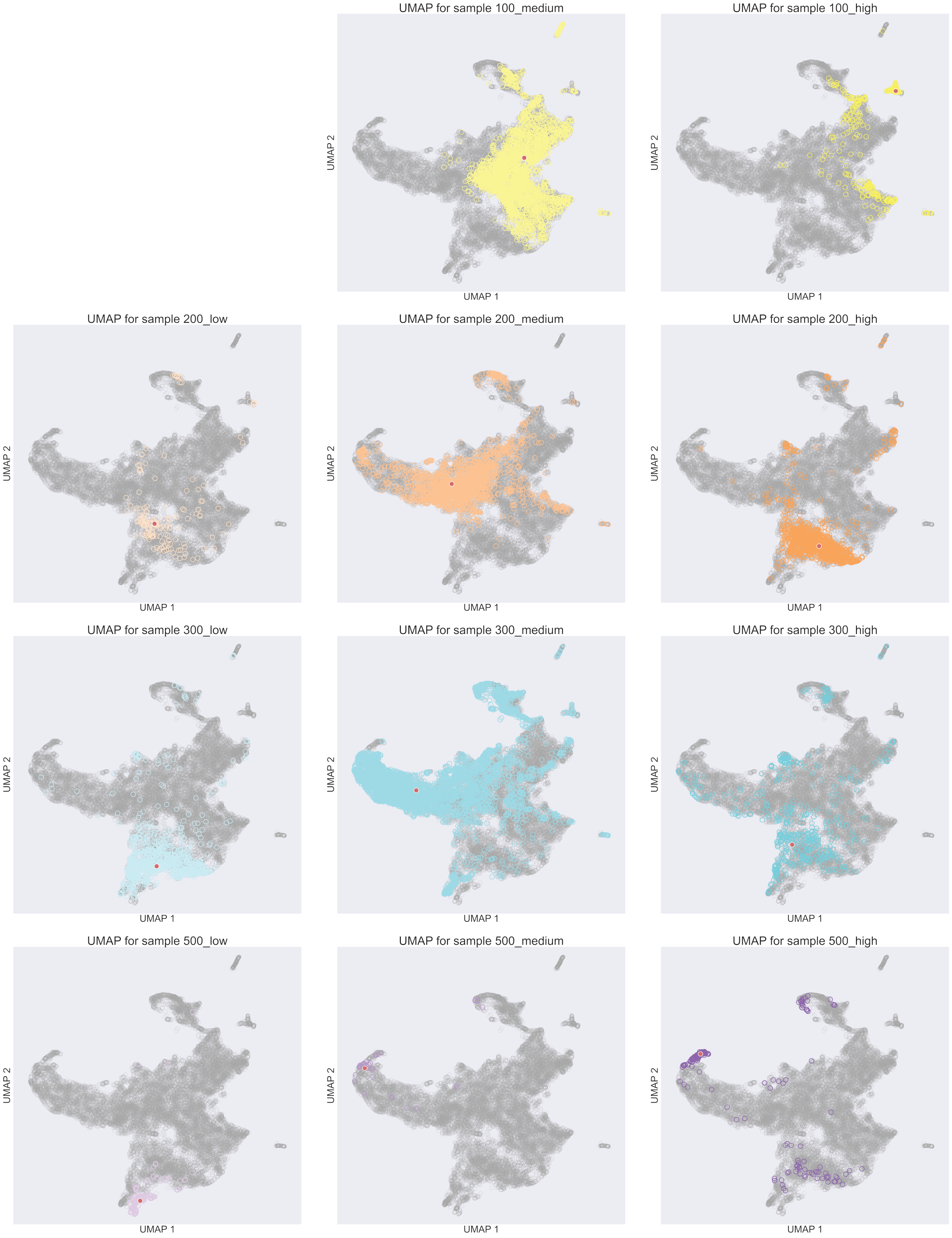


*Figure S14:* UMAP plots where the color is highlighted for each formulation in the random antibody orientation NP formulations.

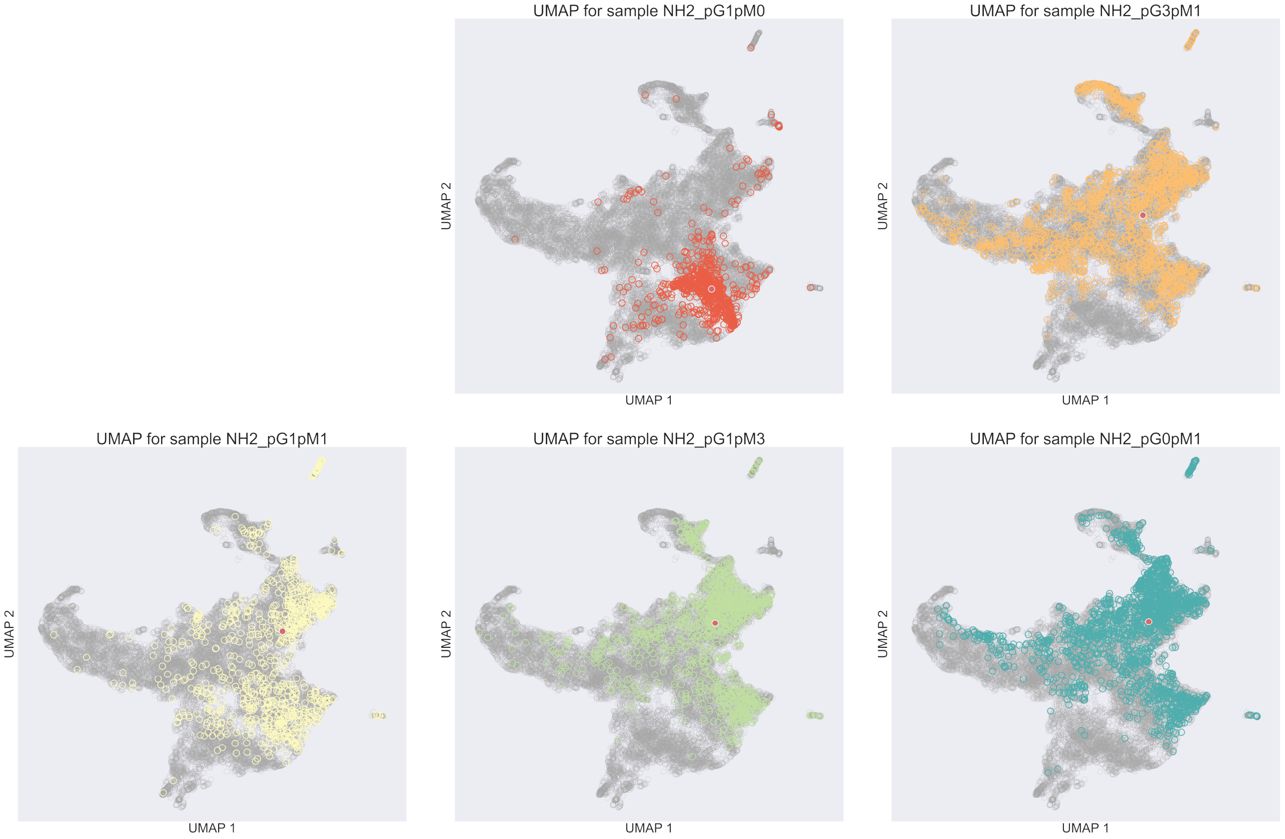


*Figure S15:* UMAP plots where the color is highlighted for each formulation in the controlled antibody orientation NP formulations.

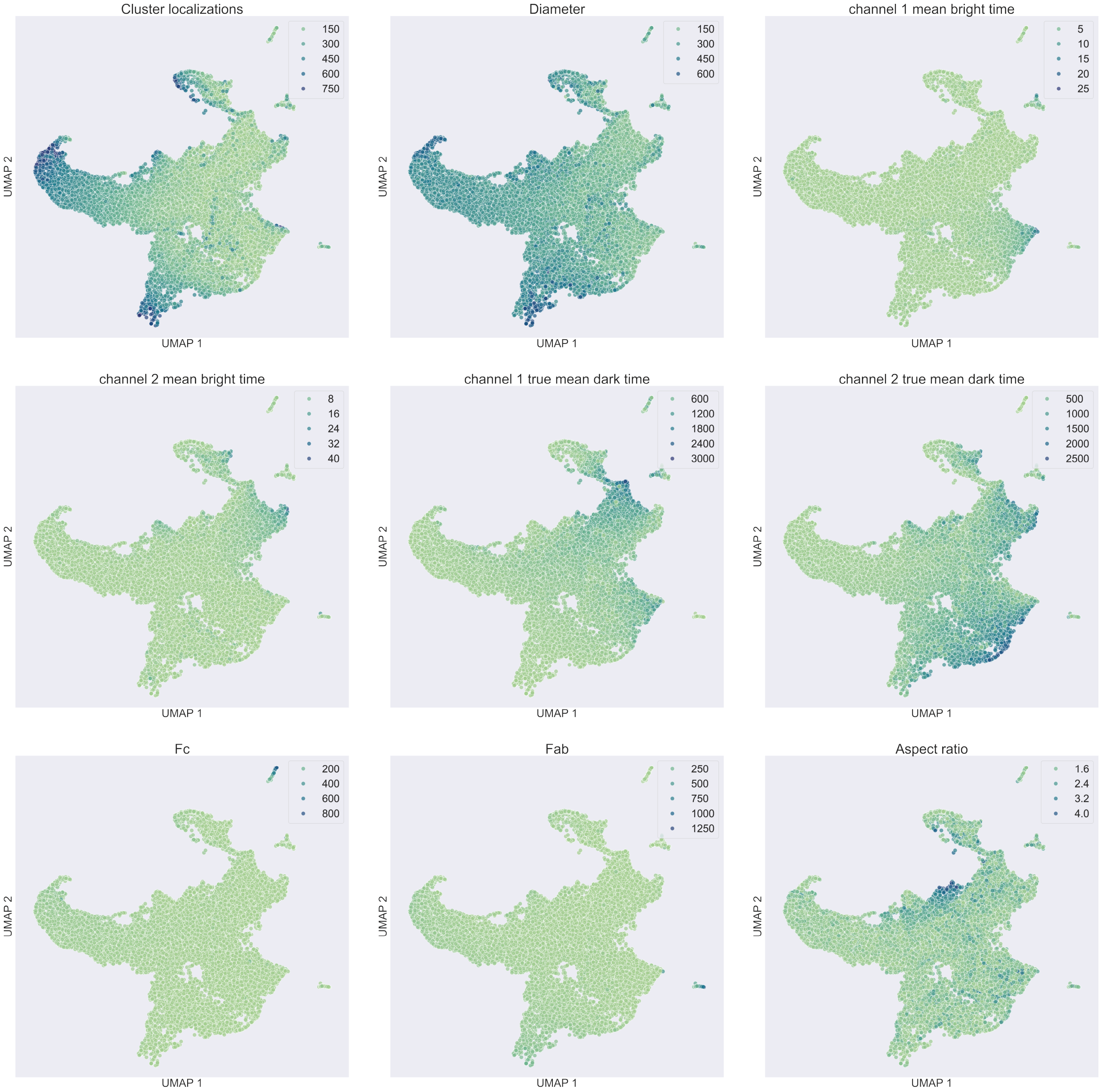


*Figure S16:* UMAP plots colored based on the values of various features. Thus, one can observe the influence of such features in the data separation.

In the case of the MST, we have separated the plots based on the experimental nature of the formulations: (a) not controlling the orientation of the antibodies (Supplementary Figure S17), and (b) controlling the antibody orientation (Supplementary Figure S18).

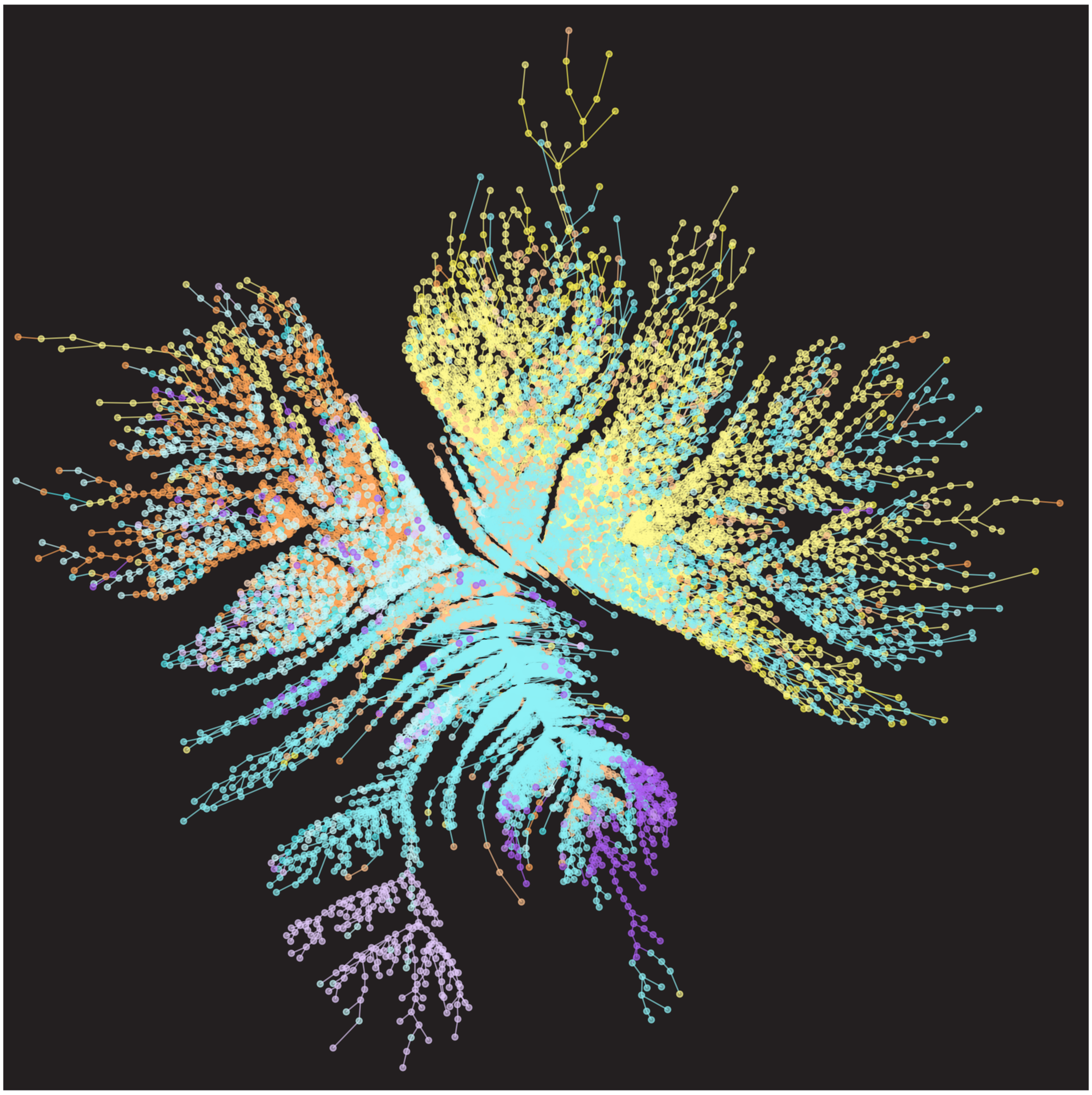

*Figure S17:* Minimum Spanning Tree (MST) of the random antibody orientation NP formulations.

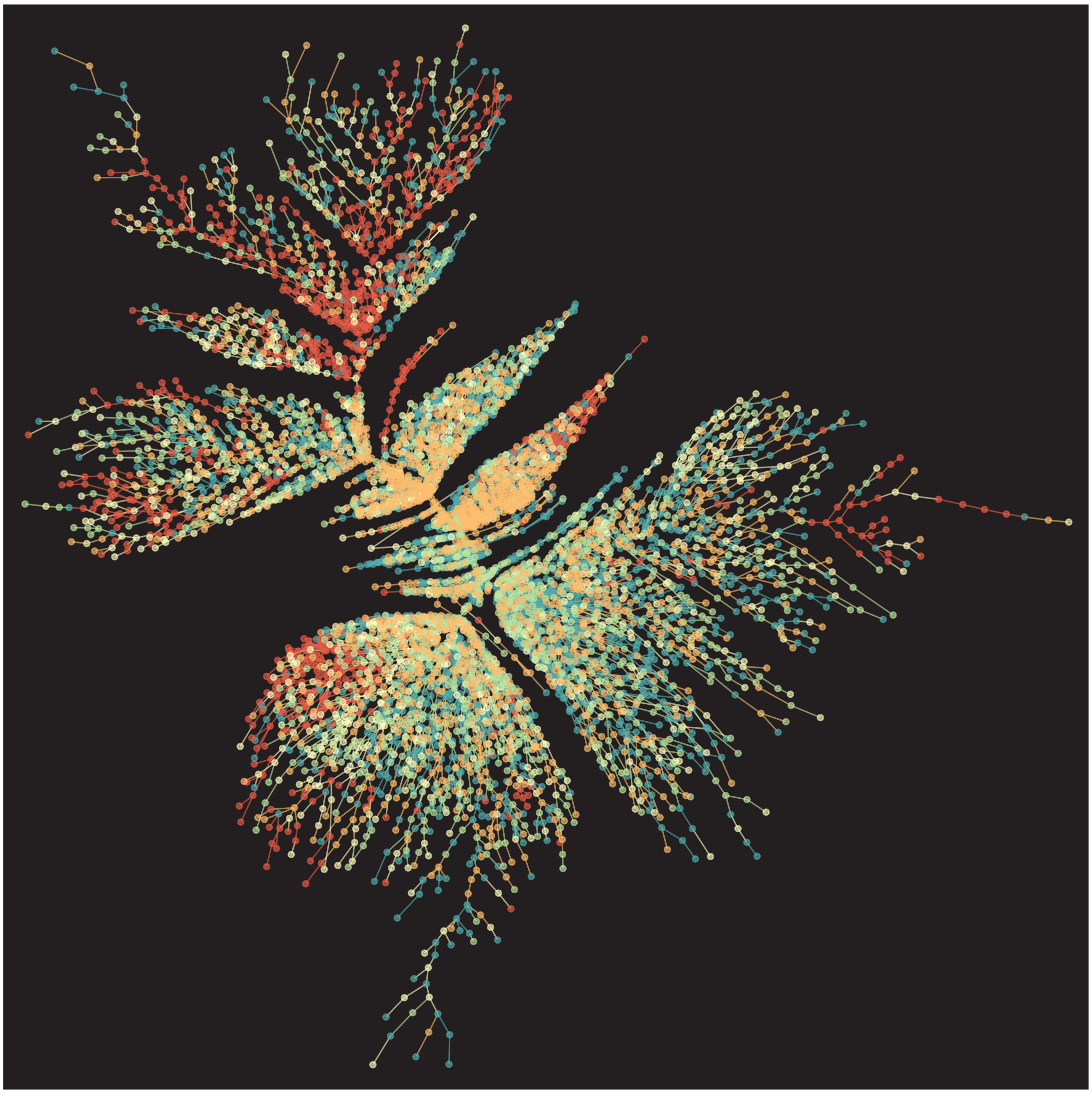

*Figure S18:* Minimum Spanning Tree (MST) of the controlled antibody orientation NP formulations.

### 4 Reproducibility studies

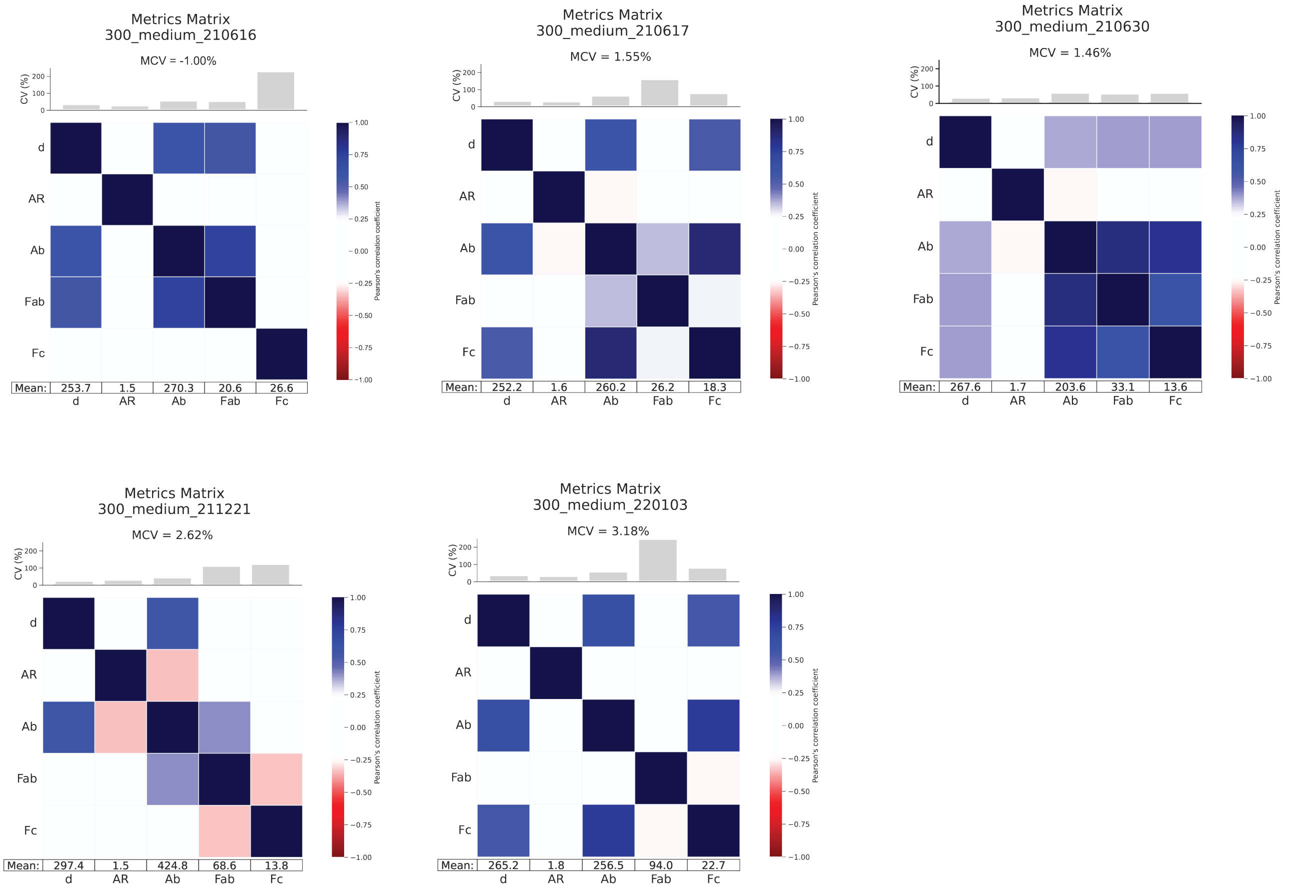


*Figure S19:* Metrics matrix for the reproducibility over time study.

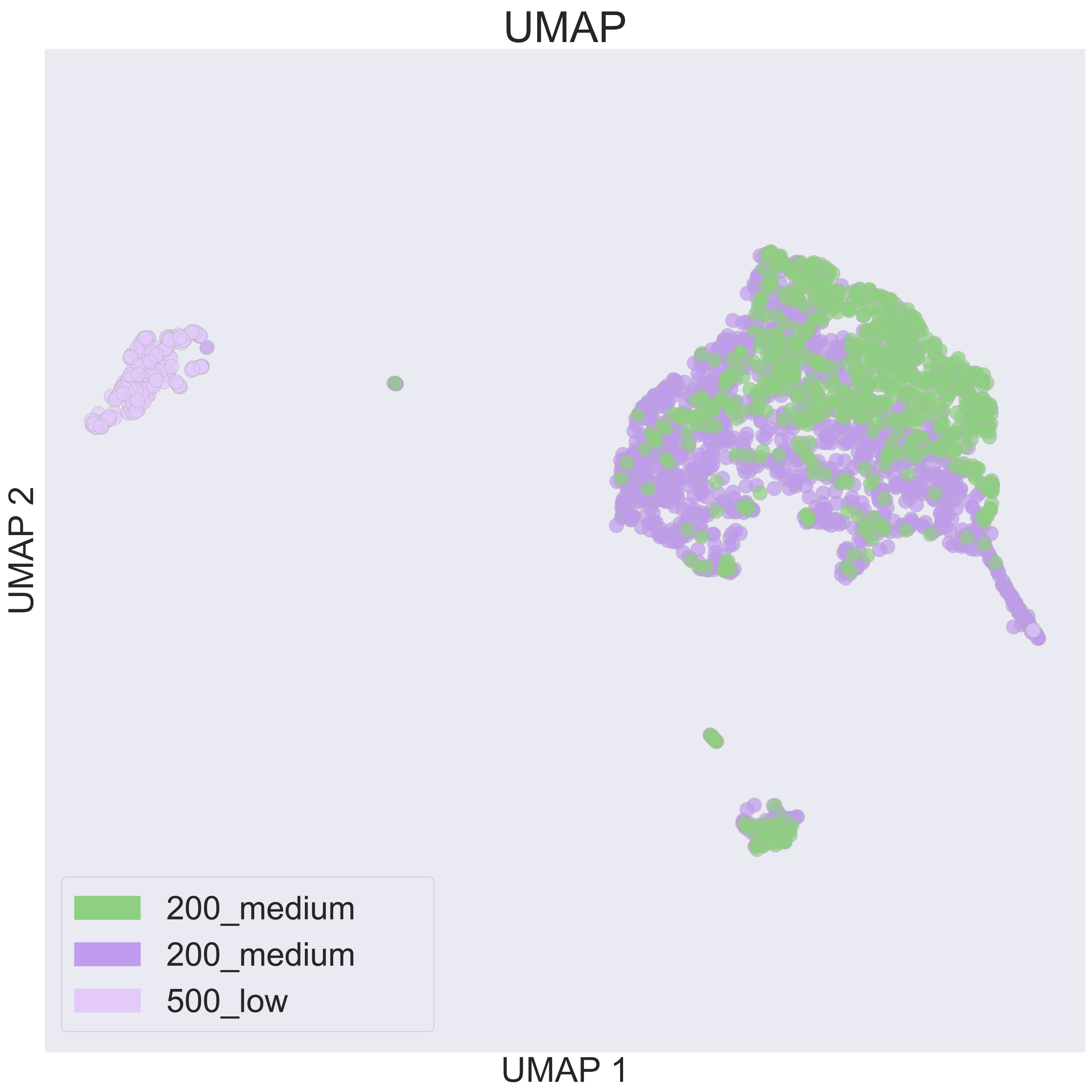


*Figure S20:* UMAP of the experiments replicated by two operators, adding an additional sample to compare the similarity between two groups of the same formulation and a different formulation.
